# Redox reactivities of membrane-bound Amyloid-β-Cu complexes and their targeting by metallothionein-3

**DOI:** 10.1101/2025.06.13.659328

**Authors:** Luciano Perez-Medina, Gabriele Meloni

## Abstract

Alzheimer’s disease (AD) is characterized by the accumulation of amyloid-β peptide (Aβ_1-40/42_) in the central nervous system (CNS) and its aggregation in senile amyloid plaques. Copper coordination to Aβ triggers Aβ_1-40/42_ aggregation and, in the presence of biological reducing agents, it promotes the catalytic generation of reactive oxygen species (ROS) via Fenton-type and Haber-Weiss reactions. Due to its amphiphilic nature, Aβ_1-40/42_ can interact with cell membranes and compromise their integrity by thinning the lipid bilayer and forming channel-like structures potentially leading to cell death. In this work, by applying biophysical and biochemical approaches, we characterized the insertion of Aβ_1-42_ into an artificial lipid bilayer system mimicking cell membranes and demonstrate that the Aβ_1-42_-lipid interaction does not prevent the Cu^2+^ coordination to Aβ_1-42_. We performed a comparative analysis of the redox reactivities of membrane-bound Aβ_1-42_ (memAβ_1-42_-Cu^2+^) species with soluble Aβ_1-42_-Cu^2+^ establishing that membrane insertion leads to memAβ_1-42_-Cu^2+^ complexes featuring an enhanced detrimental catechol oxidase activity towards the neurotransmitter dopamine. Moreover, memAβ_1-42_-Cu^2+^ efficiently catalyzes Aβ di-tyrosine crosslinking and hydroxyl radical production in the presence of ascorbate. In addition, we establish that memAβ_1-42_-Cu^2+^ redox reactivity catalyze lipid peroxidation in membranes containing polyunsaturated fatty acids (PUFAs), such as arachidonic acid (AA), leading to the generation of malondialdehyde (MDA) toxic end-products. This reactivity compromises the structural integrity of the lipid bilayers resulting in membrane leakage, further substantiating how important is to control aberrant Aβ_1-40/42_-Cu^2+^ interactions in AD.

Metallothioneins (MTs) are key metalloproteins central to neuronal and astrocytic transition metal homeostasis and buffering. These cysteine-rich proteins bind with high affinity d^10^ metals (Cu^+^ and Zn^2+^) forming two metal thiolate clusters in their N-terminal β-domain and C-terminal α-domain. The metallothionein-3 (MT-3) isoform is central to metal homeostasis in the CNS, but it is downregulated in AD patients, possessing a neuroprotective role in AD. MT-3 can control aberrant protein-Cu^2+^ interactions and the Cu-centered redox reactivities of amyloidogenic protein-Cu^2+^ complexes such as α-synuclein (Parkinson’s disease), PrP (prion disease), and soluble and aggregated Aβ_1-40_ (AD). In this work, we unravel that the detrimental memAβ_1-42_-Cu^2+^ catechol oxidase and redox reactivities can be efficiently silenced by MT-3 via metal swap reactions, effectively scavenging and reducing Cu^2+^ to Cu^+^ in its β-domain using thiolates as electron source, forming the redox-inert species Cu^+^_4_Zn^2+^_4_MT-3. Consequently, MT-3 can efficiently prevent lipid peroxidation and protect membrane structural integrity. New strategies targeting membrane-bound Aβ_1-42_-Cu^2+^ complexes as key players of the AD etiology could be envisioned.

## 1. Introduction

Alzheimer’s disease (AD) is the most diffuse neurogenerative disorder and the leading cause of dementia, a general umbrella term utilized to describe the downgrade and progressive loss of cognitive abilities.^1^ AD is a characterized by memory loss, cognitive decline, and behavioral changes correlated to distinctive pathophysiological and molecular hallmarks. Patients live with years of morbidity upon initial AD diagnosis before succumbing to the disorder. Currently 34.4-40.1 million people worldwide live with AD, and this number is expected to increase to 91.6–106.9 million by year 2050.^2^ Characteristic AD pathophysiological hallmarks include the deposition of extracellular amyloid-beta (Aβ) peptide plaques and the presence of intracellular neurofibrillary tangles composed of hyperphosphorylated tau protein in the central nervous system (CNS), leading to synaptic dysfunction and neuronal death. Additional features include diffuse oxidative stress markers and oxidative damage, neuroinflammation, brain atrophy, and impaired neurotransmitter homeostasis, particularly a decline in acetylcholine levels. No therapeutic cure for AD is currently available, and treatments are focused on slowing disease progression.

The primary molecular AD hallmark is the increased production and accumulation of the Aβ peptide, generated and released into the extracellular space upon proteolytic cleavage of the neuronal transmembrane amyloid precursor protein (APP).^3^ The β- and γ-secretases are the enzyme complexes responsible for cleaving APP, resulting in the production of the 39 to 43-residue Aβ peptide (with Aβ_1-42_ being the most diffuse and toxic peptide form in AD patients). AD belongs to protein misfolding diseases (aka amyloidosis), characterized by aberrant protein aggregation via a complex and ramified pathway that, in parallel to the generation of amorphous structures, lead to the production of characteristic insoluble protein fibrillar aggregates rich in β-sheet secondary structure.^4^ These senile plaques were first described by the German physician Alois Alzheimer.^5^ While based on the amyloid hypothesis these fibrillar plaques were considered the most toxic Aβ form,^6^ evidence suggest that small soluble globular oligomers feature a higher correlation with cognitive impairment than the larger aggregates, thus being considered the most toxic species.^7^ In agreement, large Aβ deposits in the brain of elderly people do not necessarily correlate with cognitive decline.^8^

Aβ can bind transition metal ions with high affinity at its N-terminal region (amino acids [aa] 1-16), in particular copper and zinc.^9,10^ Deposition at sub-to-mM concentrations of zinc (Zn), copper (Cu), and iron (Fe) has been demonstrated in plaques of AD patients.^10^ Transition metals coordination trigger and promote Aβ aggregation, and exacerbate its toxicity by aberrant metal-catalyzed reactivities.^11^ While copper-induced Aβ aggregation is associated with reactive oxygen species (ROS) production and neurotoxicity, zinc-induced Aβ aggregation is thought to have neuroprotective effects.^12^ ROS are generated by Aβ–Cu^2+^ through copper redox-cycling in the presence of molecular oxygen via Haber-Weiss and Fenton-like chemistry,^13^ which depends on Aβ–Cu^2+^ reduction by molecules such as ascorbate (found in neurons at mM concentration)^14^, glutathione, neurotransmitters (e.g. dopamine), and cholesterol.^9,15^ Oxidative damage and neuroinflammation are indeed hallmarks of AD brains.

At physiological pH soluble Aβ coordinates Cu^2+^ in square pyramidal geometry,^16^ and two Cu^2+^ coordination modes exist as a function of pH (**Figure 1A**). In coordination mode 1 (component 1, prominent at lower pH values), Cu^2+^ is complexed in a square pyramidal geometry by the N-terminal amino group, a carbonyl from the peptide bond between Asp1-Ala2, and two nitrogen atoms from the imidazole rings of His6 and His13/His14 (defined as component 1a if His13, or component 1b if His14) in the equatorial positions, while in the axial position a carboxylate or water molecule is weakly binding.^11,16–18^ In the coordination mode 2 (component 2) the equatorial ligands are the N-terminal amino group, the amidyl-group from the peptide bond connecting Asp1-Ala2, the carbonyl from the peptide bond between Ala2-Glu3, and an imidazole nitrogen from of any of the His residues. The transition from component 1 to component 2 occurs at pH ∼ 7.8.^16–18^ Upon Cu^2+^ reduction to Cu^+^, a shift to linear coordination occurs to two imidazole nitrogens of the His residues (**Figure 1B**). The major coordinating ligands are His13 and His14, but His6 with His13/His14 can also be observed in _Aβ-Cu+.11,16,17_

**Figure 1.**
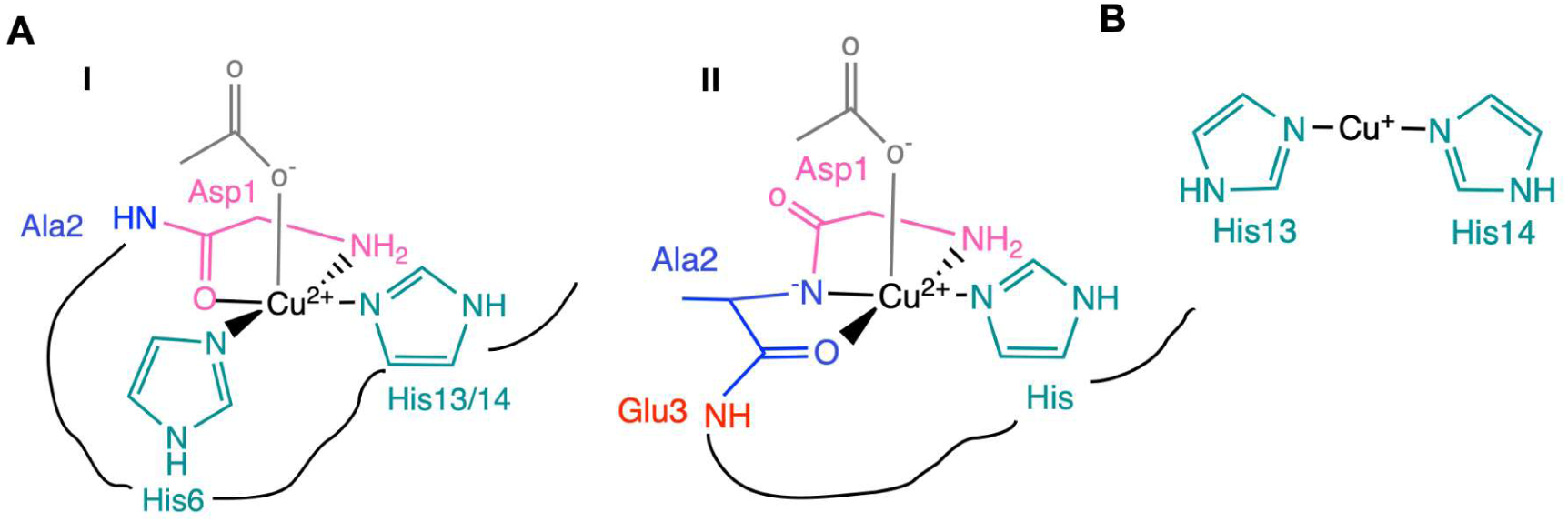
Aβ-copper coordination complexes. A) Coordination environment of Aβ-Cu^2+^ at physiological pH, with Component I and Component II highlighted. B) Major Aβ-Cu^+^ coordination complex at physiological pH.

Another detrimental interaction, significantly less investigated, is the one occurring between Aβ and lipid bilayers in biological membranes. The Aβ peptide can efficiently bind the cell membranes promoting two deleterious outcomes: 1) the formation of ion-channel-like structures that can lead to ion leakage and depolarization, and 2) the structural compression the membrane, causing lipid bilayer thinning and contributing to cellular toxicity.^19^ Altered membrane structural integrity could lead to abnormal cellular influxes of Ca^2+^ leading to mitochondrial oxidative stress and cellular toxicity.^20,21^ The aberrant redox reactivity catalyzed by Aβ-Cu^2+^ complexes with molecular oxygen, and investigations on the cellular toxicity Aβ-Cu^2+^ have been primarily focused on the soluble and aggregated species.^22–26^ However, limited information is available on the nature of Aβ-Cu^2+^ complexes generated when Aβ inserts in lipid bilayers, nor how the aberrant Aβ-Cu^2+^ reactivity is modulated upon membrane binding. Furthermore, while the reduction of aberrant Aβ-metal interactions by metal-protein attenuating compounds (MPACs) and the attenuation of oxidative stress have been investigated as potential therapeutic interventions in AD, it remains unclear how protective endogenous metal chelators can target aberrant Cu^2+^ binding in these membrane-bound complexes. Studies on other misfolding proteins-Cu^2+^ complexes relevant to other neurodegenerative disorders, such as α-synuclein (α-Syn, involved in the development of Parkinson’s disease), have demonstrated that binding to membranes can alter the redox activity and toxicity of the α-Syn-Cu^2+^ complexes when compared to the soluble α-Syn-Cu^2+^ counterparts.^27,28^

In the CNS, transition metal concentrations are tightly regulated to meet cellular requirements while avoiding metal-dependent toxicity. Together with the activity of transmembrane metal transporters and metallochaperones,^29,30^ metallothioneins (MTs) are central in controlling the metabolism of Zn and Cu. MTs are low-molecular weight proteins (61-68 aa) characterized by high Cys content and capable of binding with high affinity d^10^ transition metals, such as Zn^2+^ and Cu^+^, in stable metal-thiolate clusters.^31^ Among the four human metallothionein isoforms (MT-1/-4), the neuronal Zn_7_-metallothionein-3 (Zn_7_MT-3, also known as Growth Inhibitory Factor — GIF) is efficiently secreted in the extracellular space and play a crucial role in neutralizing and silencing the redox activity of soluble monomeric and insoluble fibrillar Aβ-Cu^2+^ complexes.^25,26^ MT-3 is abundantly expressed in neurons, and to a lesser extent astrocytes, where it plays important roles in ions Cu^+^ and Zn^2+^ homeostasis and controlling neuronal outgrowth.^32,33^ MT-3 expression is downregulated in AD patients by approximately 30%,^34^ and consequently a reduction of its unique antioxidant capacity contribute to disease progression. Zn_7_MT-3 features two Zn^2+^-thiolate clusters — a Zn^2+^_3_CysS_9_ core in the N-terminal β-domain and a Zn^2+^_4_CysS_11_ cluster in the C-terminal α-domain — with metals coordinated by 20 conserved cysteine residues.^35^ It has been shown that MT-3 efficiently controls abnormal proteins-Cu^2+^ interactions (including Aβ-Cu^2+^) via metal exchange reactions, where Zn_7_MT-3 can efficiently reduce Cu^2+^ to Cu^+^ and bind it within an oxygen-inert redox-inactive Cu^+^_4_-thiolate cluster containing two disulfide bonds in its β-domain, releasing zinc in the process.^25,26^ This metal swap reaction proceeds thorough the formation of unique long-lived and oxygen-insensitive disulfide radical anion intermediates in the β-domain,^36^ from which Zn^2+^ is released and where the thiolates provide the electron source necessary to reduce the Cu^2+^ to Cu^+^. As the β-domain of MT-3 exhibits a significantly higher binding affinity for Cu^+^ (K_D_ = 10⁻^20^ M) than for Zn^2+^, MT-3 can efficiently target high-affinity protein-Cu^2+^ complexes, including Aβ-Cu^2+^, α-Syn-Cu^2+^, and PrP-Cu^2+^, thereby preventing their copper-dependent reactivities and toxic effects.^37,38^

Due to the importance of how the reactivity of Aβ-Cu^2+^ is affected upon binding and insertion into artificial lipid bilayers, in this study we examined the metal-binding and redox properties of physiologically relevant membrane-bound Aβ-Cu^2+^ complexes and subsequently explored the capability of Zn_7_MT-3 to target and suppress their redox reactivity. We demonstrate that the catechol oxidase activity of the membrane-bound species (memAβ-Cu^2+^) against the neurotransmitter dopamine is significantly enhanced upon membrane insertion, as well establish the catalytic generation of ROS (e.g. hydroxyl radical) and di-tyrosine memAβ cross-linking in the presence of molecular oxygen and ascorbate. We also developed a platform to demonstrate that memAβ-Cu^2+^ dramatically exacerbate lipid peroxidation leading to membrane leakage upon disruption of the lipid bilayer structural integrity. This reactivity provides a molecular understanding and connection between Aβ-Cu^2+^ reactivity and membrane-related cellular toxicity.

Additionally, we show that Zn_7_MT-3 effectively target Cu^2+^ not only in soluble Aβ-Cu^2+^ but also membrane-bound Aβ-Cu^2+^ complexes, inhibiting all the redox reactions and exerting a protective role towards lipid peroxidation through unique metal-swap reactions.

## 2. Material and Methods

### 2.1. Chemicals and reagents

All reagents utilized in this work were of the highest commercially available purity. For the entire work, lyophilized Aβ_1-42_ from Anaspec Inc. (AS-20276) was utilized. The identity and purity of Aβ_1-42_ batches were verified by Matrix Assisted Laser Desorption Mass Spectrometry (MALDI-MS) or HPLC.

### 2.2. Preparation of Aβ_1-42_ stock solutions

Lyophilized Aβ_1-42_ was resuspended in 10 mM NaOH to prevent aggregation^39,40^ under constant agitation for 15 min, and centrifuged at 17,000 x*g,* 5 min, at 4 °C, to remove any insoluble material. The Aβ_1-42_ stock concentration was determined by absorption spectroscopy (ε_280_=1490 M^−1^cm^−1^)^41^ using a NanoDrop One spectrophotometer, and Chelex 100 treated MilliQ water was added to dilute the stocks and obtain Aβ_1-42_ concentrations below 100 μM. The stocks were subsequently shock-frozen in liquid N_2_ and stored at −80 °C until further use. When utilized for further experiments, the Aβ stocks were thawed at 4 °C, buffered at pH = 7.4, and the aliquots were centrifuged at 17,000 x*g*, 4 °C, 30 min, to remove any possible aggregate prior to use.

### 2.3. Preparation of Small Unilamellar Vesicles (SUVs)

Lipid mixtures (70:30; w/w) composed of 1-palmitoyl-2-oleoyl-glycero-3-phosphocholine (POPC; Avanti Polar Lipids) and 1-palmitoyl-2-oleoyl-sn-glycero-3-phospho-(1’-rac-glycerol) (sodium salt, POPG; Avanti Polar Lipids) dissolved in chloroform were prepared by rapid mixing of lipids stocks in glass balloons to obtain a final total lipid concentration of 50 mg/ml. To generate a thin lipid film, the chloroform solvent was gently evaporated under a nitrogen gas stream, under rotation. To further remove any solvent trace, lipids were further dried overnight in a vacuum desiccator. The lipid film was then resuspended in MilliQ water, Chelex 100 resin (BioRad) was added (final Chelex 100 slurry concentration: ∼ 0.1 g/mL) and the lipid suspension was mixed on a tube rotator for 2 hours at room temperature. The mixture was centrifuged at 500 x*g* for 5 min (Thermo Scientific Sorvall ST8 centrifuge) to ensure that the Chelex 100 resin completely settled. The supernatant containing the lipid suspension was aliquoted and stored at −80 °C until further use. To generate small unillamellar vesicles (SUVs) the lipid mixture was subjected to three freeze/thaw cycles followed by sonication. The sonication conditions were optimized to a total sonication time of 5 min with alternating 30s:40s sonication:rest cycles at a 60% power (Qsonica Q500). The formation of SUVs was confirmed via Dynamic Light scattering (DLS) analysis.

### 2.4. SUV dynamic light scattering analysis

Freshly sonicated lipids were diluted to a final lipid concentration of 0.2 mg/ml in 20 mM N-ethylmorpholine pH 7.4, 100 mM NaCl, or 20 mM phosphate buffer pH 7.4, and the prepared SUVs were subjected to dynamic light scattering (DLS) analysis at room temperature to determine their size distribution. The samples were placed into a polystyrene cuvette and DLS measurements performed in a Zeta sizer Nano ZS (Malvern Panalytical) utilizing the following parameters: 175° (scattering angle), 633 nm (laser wavelength), a refractive index of 1.51 and medium index of 1.33.

### 2.5. Preparation of Membrane-bound Aβ_1-42_ (memAβ)

Aβ_1-42_ stocks and the 70:30 POPC:POPG (w/w) lipid mixture after sonication (SUVs) were mixed to a protein:lipid molar ratio of 1:500 (mol/mol), and incubated in either 20 mM N-ethylmorpholine pH 7.4, 100 mM NaCl, or 20 mM phosphate buffer pH 7.4, in a temperature-controlled mixer (ThermoMixer C, Eppendorf) at 25 °C, for 12 and 3 hours, respectively, under constant shaking (300 rpm). For all experiments (except for SEC analysis), the Aβ_1-42_ concentration during incubation was 12 µM while the total lipid concentration was 6 mM (Aβ_1-42_:lipid = 1:500, mol/mol).

### 2.6. Size Exclusion Chromatography analysis of Aβ_1-42_ binding to SUVs lipid bilayers

To determine the binding of Aβ_1-42_ to the lipid vesicles, size exclusion chromatography was performed. A Superdex 75 Increase 10/300 GL column connected to an ÄKTA Pure FPLC system (Cytiva) was equilibrated 20 mM N-ethylmorpholine pH 7.4, 100 mM NaCl (3 column volumes (CV)) at a flow rate of 0.5 ml/min. Three different samples, soluble Aβ_1-42_, (20 μM), Aβ_1-42_ (20 μM) incubated with SUVs (10 mM total lipids), and SUVs (10 mM total lipids) were analyzed upon injection and elution in 20 mM N-ethylmorpholine pH 7.4, 100 mM NaCl.

### 2.7. Dot Blot analysis of Aβ_1-42_ binding to SUVs lipid bilayers

After SEC analysis, the eluted fractions were collected and analyzed by immunoblotting to detect Aβ_1-42_. A Bio-Dot Apparatus (BioRad) was employed for the analysis. All buffers needed for the immunoassay were filtered (0.22 μm; except for milk for which regular filter paper was used) prior to analysis. A 0.2 μm nitrocellulose membrane (Bio-Rad) was hydrated in 20 mM Tris/HCl pH 7.4, 150 mM NaCl (TBS). The Bio-Dot apparatus was assembled, and TBS (100 µl) was added to each well prior to applying vacuum to remove any remaining solution. This washing step was repeated twice to ensure uniform membrane hydration. Aβ_1-42_, memAβ_1-42_, and SUVs from SEC analysis (200 μl) were blotted on the nitrocellulose membrane. The Bio-Dot apparatus was covered in aluminum foil and kept at 4 °C until all the samples permeated through the membrane. A 10% non-fat milk solution (w/v; 190 µl) dissolved in TBS-T (TBS supplemented with Tween-20, 0.01% v/v) was applied to each well for blocking and incubated at room temperature for 1 h. Subsequently, the solution was removed by pipetting, and four additional washing steps were conducted by applying TBS-T (190 µl) to each well. A 4 µl aliquot of anti-Aβ 6E10 antibody (Novus Biologicals, NBP2-62566, recognizing the 1-16 N-terminus of Aβ peptide) was diluted (1:1000; v/v) in 4 ml of a 3% (w/v) Bovine Serum Albumin solution (BSA, Fisher Scientific BP1600-100) in TBS-T. The antibody solution (100 µl) was applied to each well and incubated overnight at 4 °C. The following day, TBS-T (190 µl) was applied to each well and removed manually by pipetting after a 5 min incubation. The membrane was removed from the Bio-Dot apparatus and washed four times in TBS-T. A secondary antibody (2.5 µl; alkaline phosphatase conjugated anti-rabbit IgG; Sigma-Aldrich A3812) was diluted (1:10000; v/v) in 3% BSA/TBS-T (w/v; 25 ml), added to the membrane, and incubated for 1 hour at room temperature. The membrane was washed four times with TBS-T and developed using a nitro blue tetrazolium chloride and 5-bromo-4-chloro-3-indolyl phosphate tablet (Sigma-Aldrich B5655) dissolved in water. The dot blot protocol was adapted from the literature.^42^

### 2.8. Expression and purification of human metallothionein-3 (Zn_7_MT-3)

Human MT-3 was expressed in *E. coli* BL21(DE3)pLysS as a Cd metalloprotein following the method developed by Vašák.^43^ Protein expression was induced upon addition of 1 mM isopropyl β-D-1-thiogalactopyranoside, and after 30 min cadmium sulfate 0.4 mM were supplied to the cell media to express Cd-containing metallothionein-3. The protein was purified as previously described via two ethanol precipitation steps, followed by size exclusion and ion exchange chromatography. Apo MTs stocks were prepared by treating the samples with HCl, following the protocol developed by Vašák.^44^ These were subsequently reconstituted into the Zn_7_MT forms by adding ZnCl_2_ and adjusting the pH to 8.0 with 1 M Tris. A final size exclusion chromatography step was performed upon reconstitution on a Superdex 75 column connected to an Äkta FPLC system (Cytiva) and the protein eluted in 20 mM N-ethylmorpholine pH 7.4, 100 NaCl, or 20 mM phosphate buffer pH 7.4. Protein concentration was determined spectrophotometrically (Agilent Cary 300 UV-Vis Spectrophotometer) in 0.1 mM HCl, using an extinction coefficient of ε_220_ = 53,000 M^−1^ cm^−1^.^25^ The cysteine-to-protein ratio was assessed through photometric detection of sulfhydryl (CysSH) groups upon reaction with 2,2-dithiodipyridine in a 0.2 M sodium acetate pH 4.0, 1 mM EDTA buffer, using the extinction coefficient ε_343_ = 7,600 M^−1^ cm^−1^.^45^ Metal-to-protein ratios were determined by Inductively Coupled Plasma Mass Spectrometry (Agilent 7900 ICP-MS; instrument equipped with an autosampler) after sample digestion in 1% HNO_3_. For ZnMT-3, the measured CysSH-to-protein ratio was 20 ± 3, while the zinc-to-protein ratios were 7.0 ± 0.4.

### 2.9. Analysis of Aβ_1-42_ and memAβ_1-42_ dopamine oxidase activity

Soluble Aβ_1-42_ and memAβ_1-42_ samples (12 μM) in 20 mM N-ethylmorpholine pH 7.4, 100 mM NaCl were mixed with a CuCl_2_ stock (1 mM in H_2_O) to obtain a final Cu^2+^ concentration of 10 μM, and incubated for 20 min at room temperature. Samples were supplemented with 3-methyl-2-benzothiazolinone hydrazone (MBTH; 3 mM) from a 20 mM stock in H_2_O. The samples were aliquoted into 96-wells flat bottom plates (Corning Costar, 3370) and supplemented with dopamine (serial dilutions ranging 0.05-3 mM). Immediately after dopamine addition, the catechol oxidation was monitored by quantifying the formation of the adduct between MBTH and dopaquinone (ε_500_ = 32,500 M^−1^ cm^−1^) at room temperature. The absorbance at 500 nm was recorded every minute for 2 hours using a plate reader (Agilent BioTek Synergy H4 Hybrid Microplate Reader). The amount of dopamine generated after 85 min was utilized to fit the catalytic parameters with a Michaelis-Menten equation. Data were corrected by subtracting the recorded contribution of dopamine auto-oxidation measured in the absence of Aβ_1-42_ and memAβ_1-42_.

To test the quenching effect of Zn_7_MT-3 towards the Aβ catechol oxidase activity, Aβ_1-42_ and memAβ_1-42_ (12 μM) in 20 mM N-ethylmorpholine pH 7.4, 100 mM NaCl, were supplemented with 10 μM Cu^2+^ (to generate soluble Aβ_1-42_-Cu^2+^ and memAβ_1-42_-Cu^2+^), 3 mM MBTH, and 2 mM dopamine. The dopamine oxidation reaction was recorded in the absence and presence of Zn_7_MT-3 (2.5 μM) upon adding Zn_7_MT-3 stocks in 20 mM N-ethylmorpholine pH 7.4, 100 mM NaCl, and incubating the samples for 1 hour at room temperature prior to the addition of MBTH and dopamine. Absorbance spectra (450-650 nm) were recorded every 5 min for 2 hours (Agilent Cary 300 UV–Vis Spectrophotometer). The contribution of dopamine auto-oxidation was subtracted prior to data analysis.

### 2.10. Effect of H_2_O_2_ on the Aβ_1-42_ and memAβ_1-42_ dopamine oxidase activity

Soluble Aβ_1-42_ and memAβ_1-42_ (12 μM) in 20 mM N-ethylmorpholine pH 7.4, 100 mM NaCl, were incubated with CuCl_2_ (10 µM) for 20 min at room temperature. The effect of H_2_O_2_ on the catechol oxidase activity of Aβ_1-42_-Cu^2+^ and memAβ_1-42_-Cu^2+^ at different concentrations of dopamine (0.05 to 3 mM) was tested upon addition of MBTH (3 mM). Samples were supplemented with H_2_O_2_ (0-50 mM) using a 0.5 M stock in H_2_O. Samples were placed in 96-wells flat bottom plates (Corning Costar, 3370) and the absorbance at 500 nm (MBTH-dopaquinone adduct) was recorded every minute for 2 hours at room temperature using a plate reader (Agilent BioTek Synergy H4 Hybrid Microplate Reader). The data generated were subjected to Michaelis-Menten analysis as a function of dopamine substrate concentration for each H_2_O_2_ concentration (dopaquinone quantified at 85 min from the start of the reaction). The obtained V_max_ were subsequently plotted as a function H_2_O_2_ concentration for Michaelis-Menten analysis.

### 2.11. Determination of di-tyrosine crosslinking by memAβ_1-42_-Cu^2+^

Soluble Aβ_1-42_ and memAβ_1-42_ (12 μM) in 20 mM phosphate buffer pH 7.4, were mixed with 10 μM Cu^2+^ to generate Aβ_1-42_-Cu^2+^ and memAβ_1-42_-Cu^2+^. Samples were placed in a black wall-quartz cuvette in a Horiba FluoroMax-4 spectrofluorometer with a controlled temperature of 37 °C. A 20 mM ascorbate stock solution was equilibrated at the same temperature in a thermomixer, and subsequently added to Aβ_1-42_-Cu^2+^ and memAβ_1-42_-Cu^2+^ at a final concentration of 3 mM. The generation of the di-tyrosine crosslinked products was monitored by recording the fluorescence emission spectrum between 390 nm and 520 nm (λ_ex_ = 325 nm; slit width = 10 nm) for 30 min. To comparatively quantify the amount of di-tyrosine generated, single fluorescence emission data at λ_em_ = 425 nm were collected and analyzed as a function of time. Similar experiments were conducted upon adding 0.25 eq. of Zn_7_MT-3 to Aβ_1-42_-Cu^2+^ and memAβ_1-42_-Cu^2+^, prior to ascorbate addition, to test the MT-3 redox-silencing abilities in preventing di-tyrosine crosslinking.

### 2.12. Determination of hydroxyl radical generation by memAβ_1-42_-Cu^2+^

To investigate the hydroxyl radical production, soluble Aβ_1-42_ and memAβ_1-42_ (12 μM) in 20 mM phosphate buffer pH 7.4, were mixed with 10 μM Cu^2+^ to generate Aβ_1-42_-Cu^2+^ and memAβ_1-42_-Cu^2^. Aβ_1-42_-Cu^2+^ and memAβ_1-42_-Cu^2+^ were subsequently mixed with 1.2 mM ascorbate and 800 μM coumarin-3-carboxylic acid (3-CCA). The generation of hydroxyl radical was followed by monitoring the fluorescence emission of the 7-OH-3-CCA product at 450 nm (λ_ex_ = 395 nm; slit width = 5 nm) for 2000 s. To test the redox silencing by Zn_7_MT-3, the Aβ_1-42_-Cu^2+^ and memAβ_1-42_-Cu^2+^ complexes were incubated with 0.25 eq of Zn_7_MT-3 for 1 h at 25°C before addition of ascorbate.

### 2.13. Lipid peroxidation assays

To test whether the redox activity of memAβ_1-42_-Cu^2+^ can compromise the chemical and structural integrity of the lipid bilayers, lipid mixtures, prepared as described above, were doped with 1,2-diarachidonoyl-sn-glycero-3-phosphocholine (AALip; 1% w/w). The lipids were dried for 3-4 hours instead of overnight to minimize AALip oxidation and the total sonication time was increased to 7 min (30s:40s sonication:rest cycles with a 60% power intensity; Qsonica Q500).

Lipid peroxidation was determined by quantification of the end-product malondialdehyde (MDA). After incubation of AALip (6 mM) and Aβ_1-42_ (12 µM) in 20 mM phosphate buffer pH 7.4 for 3 hours, Cu^2+^ was added (10 µM). The samples were incubated for 20 hours at 37 °C, upon addition of 1 mM of ascorbate (from a 100 mM stock in H_2_O) every hour in the first 6 hours and further additions in the last 4 hours of incubation. The samples were subsequently centrifuged at 500 x*g* for 3 min, 4 °C. MDA was quantified using an MDA Colorimetric Assay Kit (Invitrogen, EEA015). The kit acid reagent solution was added to the samples, followed by the chromogenic agent solution. The tubes were perforated with a small syringe and incubated for 40 min at 100 °C. The samples were subsequently placed on ice for a few seconds and centrifugated at 9,600 x*g*, 20 min, 4 °C. The supernatant was recovered and placed in 96-wells plates. The absorbance of the MDA-TBA adduct at 532 nm was recorded using a Tecan Spark 20 plate reader, and the concentration of MDA was determined using a calibration curve using a MDA standard solution. To determine redox silencing properties of Zn_7_MT-3, similar experiments were conducted upon incubating memAβ_1-42_-Cu^2+^ with 0.25 eq of Zn_7_MT-3 for 1 h at 25°C before the first addition of ascorbate. Samples containing lipids but no ascorbate were used as blank, and their contributions were subtracted from all the samples.

### 2.14. Membrane leakage assays

To determine membrane leakage upon lipid peroxidation, 5(6)-carboxyfluorescein (100 μM) was encapsulated into the AALip mixture by performing six freeze/thaw cycles in liquid nitrogen, followed by sonication (30s:40s sonication:rest cycles, with a 60% power for 7 min). The samples were centrifuged at 190,000 x*g*, 2 hours, 4 °C (Thermo Scientific Sorvall MX-120 Plus Micro-Ultracentrifuge). The supernatant was discarded, and the pellet was washed in buffer (20 mM phosphate pH 7.4) and further collected by ultracentrifugation. The vesicle pellet was resuspended to the desired volume (0.5-1 mL) and incubated at 25 °C for 3 hours with Aβ_1-42_ under constant agitation (300 rpm). After the addition of Cu^2+^ (10 µM), the samples were incubated at 37 °C upon adding 1 mM ascorbate every hour for a total of 6 hours. The samples were mixed (300 rpm) for an additional 6 hours. The vesicles were pelleted via centrifugation at 190,000 xg, 2 hours, 4 °C (Thermo Scientific Sorvall MX-120 Plus Micro-Ultracentrifuge), and the 5(6)-carboxyfluorescein fluorescence emission spectrum of the supernatant was recorded from 500 to 650 nm (λ_ex_ = 450 nm; slith width = 10 nm) using a Tecan Spark 20 plate reader. The pellet was resuspended in buffer and Triton-X 100 was added to a final concentration of 2% (v/v). After 30 min, the sample was centrifuged at 190,000 xg, 2 hours, 4 °C (Thermo Scientific Sorvall MX-120 Plus Micro-Ultracentrifuge), and the fluorescence emission was recorded. Samples containing lipids but no ascorbate were used as blank and their contributions were subtracted from all the samples. To test the capability of Zn_7_MT-3 in protecting from lipid peroxidation and membrane leakage, similar experiments were conducted upon incubating memAβ_1-42_-Cu^2+^ with 0.25 eq of Zn_7_MT-3 for 1 h at 25°C before the first addition of ascorbate.

### 2.15. Low-temperature (77K) luminescence characterization of the metal swap reaction between Zn_7_MT-3 and Aβ-Cu^2+^ complexes

MemAβ-Cu^2+^ ([Aβ_1-42_] = 12 µM and [Cu^2+^] = 10 µM) and Zn_7_MT-3 (2.5 µM) were mixed and incubated for 1 hour at room temperature. The samples were centrifuged at 190,000 x*g,* 2 hours, 4 °C (Thermo Scientific Sorvall MX-120 Plus Micro-Ultracentrifuge). The supernatant was recovered, and the pellet was resuspended in 20 mM N-ethylmorpholine pH 7.4, 100 mM NaCl, to the original volume, to be analyzed by low temperature luminescence, SDS-PAGE, silver staining, and ICP-MS. Low-temperature luminescence emission spectra and lifetime decays were collected using a FluoroMax-4 spectrofluorometer (Horiba Scientific). The samples were transferred into quartz tubes with a 2 mm inner diameter, frozen in liquid N_2_, and placed in a cylindrical quartz Dewar filled with liquid N_2_. Emission spectra (380–750 nm, slit width = 5 nm) were recorded at 77 K using an excitation wavelength of 320 nm (slit width = 5 nm), with an initial delay of 10 μs and a sample window of 300 μs. Lifetime measurements were conducted for the emission bands at 425 nm and 575 nm, employing an initial delay of 50 μs and a 300 μs sample window. Delay increments of 10 μs and 20 μs, along with maximum delays of 500 μs and 1000 μs, were applied for the 425 nm and 575 nm bands, respectively. Emission lifetimes were determined by fitting the data with a single exponential decay function.

### 2.16. Metal content determination by ICP-MS

MemAβ_1-42_ and 12 µM Aβ_1-42_ (12 µM) were incubated with 10 µM CuCl_2_ for 20 min and subsequently subjected to ultracentrifugation (190,000 x*g,* 2 hours, 4 °C; Thermo Scientific Sorvall MX-120 Plus Micro-Ultracentrifuge). The resuspended pellets and supernatants from each sample were digested overnight in 11.5% HNO_3_ (v/v; metal-free grade) at 85 °C using a Thermomixer (Eppendorf). The following day, the samples were centrifuged (4000 x*g*, 5 min, 4 °C) and diluted with Milli Q water to obtain a final HNO_3_ concentration of 3% (v/v). The metal content was determined on an Inductively Coupled Plasma Mass Spectrometer (Agilent 7900) equipped with an autosampler.

The samples upon metal swap reactions between memAβ-Cu^2+^ ([Aβ_1-42_] = 12 µM and [Cu^2+^] = 10 µM) and Zn_7_MT-3 (2.5 µM) obtained for the low-temperature luminescence measurements, were separated by ultracentrifugation, and samples for ICP-MS prepared as described above.

### 2.17. SDS-PAGE analysis and silver staining

MemAβ_1-42_ (12 µM) and Aβ_1-42_ (12) µM were incubated for 20 min with 10 µM CuCl_2_ and then, they were separated by ultracentrifugation (190,000 x*g,* 2 hours, 4 °C; Thermo Scientific Sorvall MX-120 Plus Micro-Ultracentrifuge). The supernatants and resuspended pellets were diluted in a 3:1 ratio with Laemmli buffer containing 10% β-mercaptoethanol (Biorad; v/v). Samples were loaded on a pre-casted 4–15% polyacrylamide gel (Biorad) to perform SDS-PAGE analysis (160 V, 30 min). The gel with the Aβ_1-42_ samples was stained with 3% (w/v) Coomassie blue solution. Meanwhile, the gel for the memAβ_1-42_ was washed two times with Milli Q water and samples were fixed (15 min, twice) with fixing solution (30% ethanol:10% acetic acid:60% water). After washing the gel with 10% (v/v) ethanol twice for 5 min, gels were quickly rinsed in H_2_O to remove any ethanol trace. Using a Pierce Silver Stain Kit (Thermo Scientific), the gel was incubated with the sensitizer solution for 1 min and subsequently washed in H_2_O. The staining and enhancer solutions were mixed, and the gel incubated for 30 min in the mixture. Upon washing the gel water, the developer and enhancer solutions were mixed to silver stain the gel. Once the protein bands appeared (approx. 2-3 min after addition), the reaction was quenched with a 5% solution of acetic acid (v/v) and the gels imaged with a ChemiDoc Touch Imaging System (Biorad).

Upon the reaction between Zn_7_MT-3 (2.5 µM) and memAβ_1-42_ -Cu^2+^ ([Aβ_1-42_] = 12 µM and [Cu^2+^] = 10 µM), as described for the low temperature luminescence characterization, the samples were subjected to ultracentrifugation (190,000 x*g,* 2 hours, 4 °C), and the supernatants and resuspended pellets were diluted in a 3:1 ratio with Laemmli buffer containing 10% β-mercaptoethanol (Biorad; v/v). SDS-page analysis with silver staining was conducted as described above.

## 3. Results and discussion

### 3.1. Generation of membrane bound Aβ-Cu^2+^ complexes

In the CNS of AD patients, the accumulation of Aβ peptide is postulated to result in binding and insertion into biological membranes by following an off-pathway independent from that classical fibrillar aggregation into amyloid plaques. However, the redox behavior of Aβ-Cu^2+^ embedded in lipid bilayers remains poorly investigated. To assess how membrane insertion influences Aβ-Cu^2+^ redox activity in comparison to Aβ soluble forms, Aβ_1-42_ was selected as it represents the most toxic Aβ form and it is the Aβ species most prone towards aggregation and membrane binding.

To investigate Aβ_1-42_ interactions with lipid membranes, we assessed the incorporation soluble Aβ_1-42_ into small unilamellar vesicles (SUVs) composed of a phosphatidylcholine (POPC, zwitterionic) and phosphatidylglycerol (POPG, negatively charged) mixture at a 70:30 (w/w) ratio, generated by sonication. This lipid composition was demonstrated to minimize Cu^2+^ interactions with lipid head groups.^46^ The generation of SUVs was confirmed by dynamic light scattering (DLS). DLS analysis showed average vesicles size of approx. 50 nm with a satisfactory polydispersity index of 0.270 (**Figure 2A**). To generate membrane bound Aβ_1-42_ complexes a protein-to-lipid molar ratio of 1:500 (mol/mol) was selected, a ratio demonstrated to promote binding of other amyloidogenic proteins to membranes (e.g. α-Syn-Cu).^27,46^ Aβ stocks solutions were incubated in N-ethylmorpholine pH 7.4 buffer to promote the insertion of the peptide into the artificial lipid bilayer. Complete insertion of Aβ_1-42_ into lipid bilayers was confirmed by independent ultracentrifugation and size-exclusion chromatography analysis, followed by SDS-PAGE and dot-blot, respectively. Upon Aβ_1-42_ incubation with SUVs, lipid vesicles were separated by ultracentrifugation and the resuspended pellet and supernatant were analyzed by SDS-PAGE followed by silver staining. In agreement with complete Aβ_1-42_ insertion in the lipid bilayer, a ∼ 4kD band corresponding to Aβ_1-42_ was exclusively observed in the pellet fraction but not in the supernatant (**Figure 2B**). Complete binding of Aβ_1-42_ to lipid bilayers was also independently confirmed by size-exclusion chromatography. Aβ_1-42_ in the absence of lipids, Aβ_1-42_ incubated with SUVs, and lipid vesicles samples were injected into a Superdex 75 column followed by an immuno dot-blot assays using an anti-Aβ antibody (6E10) to identify the presence of Aβ_1-42_ in the collected fractions (**Figure 2 C-D**). While elution of Aβ_1-42_ without lipids was exclusively detected in the peak at 15 mL (corresponding to soluble Aβ_1-42_), in the presence of SUVs Aβ_1-42_ coeluted exclusively at the column void volume (∼ 8.2 mL), where SUVs elute. Peak integration and dot blot analysis corroborated Aβ_1-42_ insertion in lipid bilayers.

**Figure 2.**
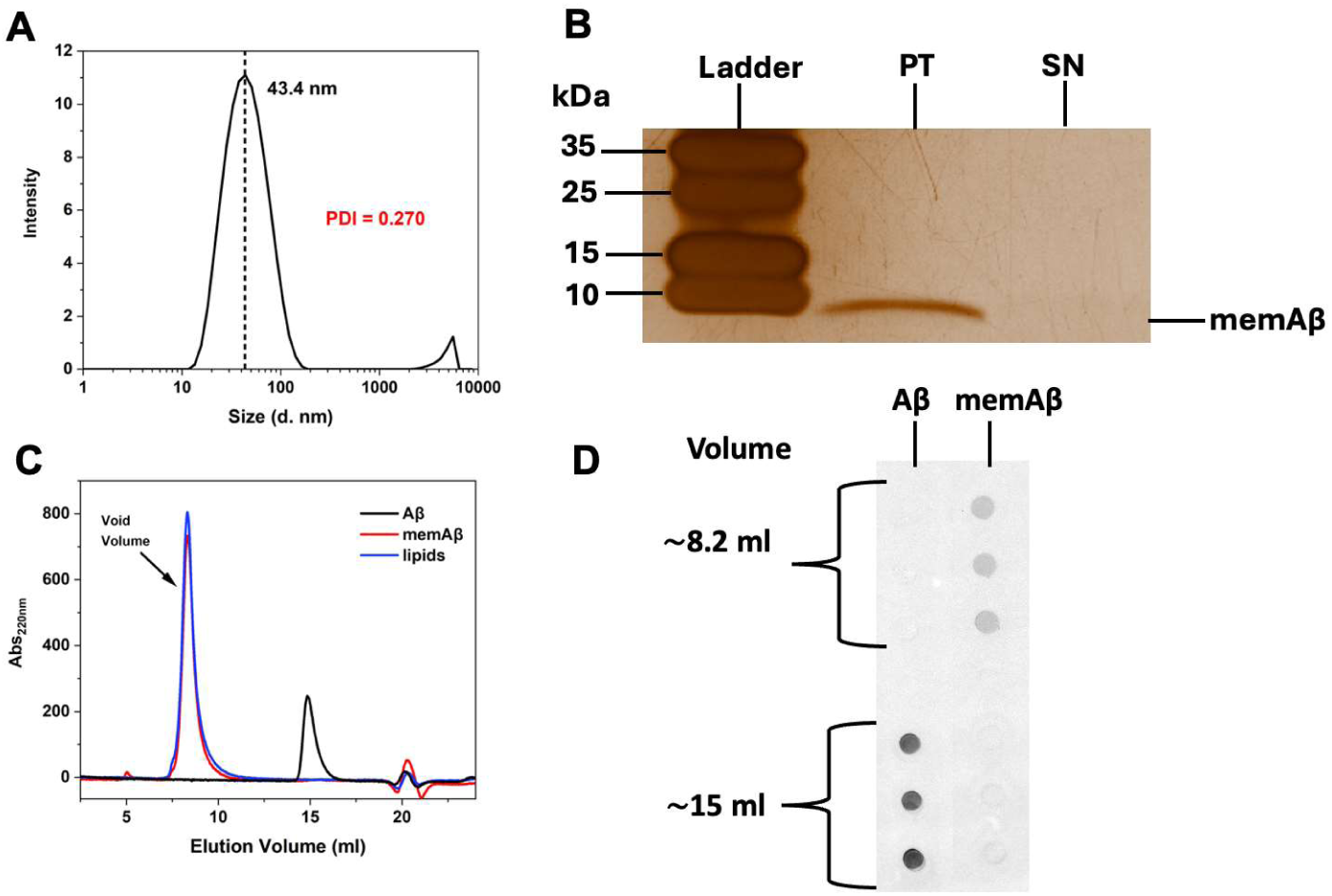
Characterization of Aβ_1-42_ insertion in Small Unilamellar Vesicles (SUVs). A) Dynamic light scattering analysis of a POPC:POPG lipid mixture (0.2 mg/ml) after sonication confirming the formation of Small Unilamellar Vesicles (SUVs) with a satisfactory polydispersity index. B) SDS-PAGE analysis and silver staining of memAβ_1-42_ SUVs upon ultracentrifugation, with pellet (PT, SUVs) and supernatant (SN) fractions, revealing the presence of Aβ_1-42_ exclusively in the PT fraction. C) Size exclusion chromatogram of soluble Aβ (20 μM; black), Aβ incubated with 10 mM lipids (memAβ_1-42_ ; red), and 10 mM lipids (blue) in 20 mM N-ethylmorpholine pH = 7.4, 100 mM NaCl. The memAβ_1-42_ and lipid SUVs eluted at the void volume, while soluble Aβ_1-42_ eluted within the separation range of the column. D) Dot blot immunoassay analysis of the SEC fractions using an anti-Aβ antibody (6E10). Soluble Aβ_1-42_ species eluted around 15 mL and memAβ_1-42_ at 8.2 mL.

To confirm that insertion of Aβ_1-42_ into the membrane does not affect its Cu^2+^ binding ability, we titrated Cu^2+^ into membrane-bound Aβ_1-42_ (at a 1.2:1 Aβ_1-42_:metal molar ratio) and quantified copper content in the lipid pellet and soluble fraction by Inductively Couple Plasma Mass Spectrometry (ICP-MS; **Figure 3A**). Upon incubation, SUVs were separated by ultracentrifugation to confirm that incubation with Cu^2+^ did not affect Aβ_1-42_ membrane binding, as demonstrated by SDS-PAGE analysis followed by silver staining (**Figure 3B**). ICP-MS analysis confirmed that all copper was recovered in the pellet fraction where membrane-bound Aβ_1-42_ is present, confirming formation of memAβ_1-42_-Cu^2+^, while copper and Aβ_1-42_ were recovered in the supernatant fraction in the absence of SUVs (**Figure 3A and 3C**), consistent with the formation of soluble Aβ_1-42_-Cu^2+^ in the absence of lipids, and confirming that the addition of the metal did not result in rapid Aβ_1-42_ aggregation. These analyses establish that the generated native-like artificial membranes allow the stoichiometric formation and reactivity investigation of membrane bound memAβ_1-42_-Cu^2+^ complexes.

**Figure 3.**
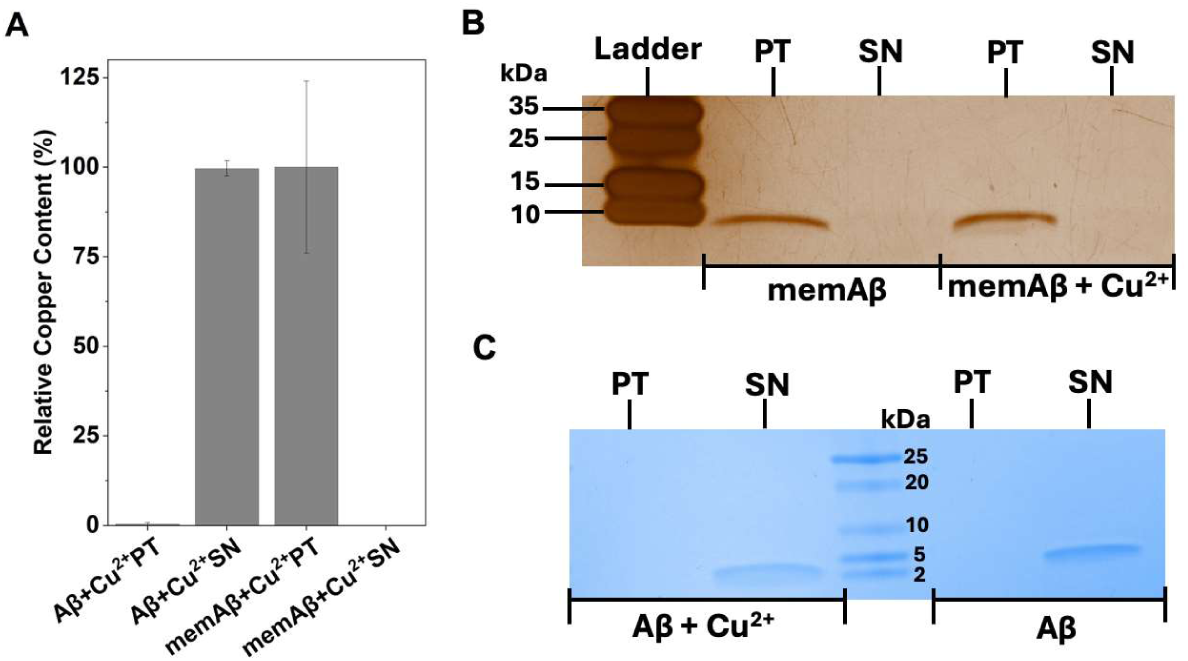
Characterization and quantification of Aβ-Cu^2+^ complexes after ultracentrifugation. A) Inductively Couple Plasma Mass Spectrometry (ICP-MS) analysis reporting relative copper content in samples obtained upon soluble Aβ_1-42_ and memAβ_1-42_ incubation with Cu^2+^. Copper is exclusively recovered in the supernatant fraction (SN) for soluble Aβ_1-42_, and in the pellet (PT) for memAβ_1-42_. Thus, the insertion of Aβ_1-42_ into the membrane does not prevent its Cu^2+^ coordination ability (n = 3 ± s.d.) B) SDS-PAGE analysis and silver staining of memAβ_1-42_ and memAβ_1-42_-Cu^2+^ samples after ultracentrifugation. C) SDS-PAGE analysis of soluble Aβ_1-42_ and Aβ_1-42_-Cu^2+^ samples upon ultracentrifugation in the absence of SUVs.

### 3.2. MemAβ-Cu^2+^ acts as a catechol oxidase catalyst and potentiate the oxidation of the endogenous neurotransmitter dopamine

It has been previously demonstrated that soluble Aβ-Cu^2+^ possess an enzyme-like phenol and catechol oxidase activity, observed with the oxidation of the 1,2,3-trihydroxylbenzene substrate (THB),^22–24^ suggesting a potential detrimental role of Aβ-Cu^2+^ in the oxidation of key neurotransmitters like dopamine in the AD brain.

We therefore examined how Aβ_1-42_ membrane insertion influences the dopamine oxidase activity compared to soluble Aβ_1-42_-Cu^2+^ complexes towards catalyzing dopamine ortho-quinone formation upon oxidation, using 3-methyl-2-benzothiazolinone hydrazone (MBTH) which forms a colored adduct with dopaquinone to monitor the reaction by absorption spectroscopy (**Figure 4A**). Absorption spectra analysis of the MBTH-adduct with dopamine ortho-quinone showed a substantial increase in ortho-quinone production for memAβ_1-42_-Cu^2+^ compared to Aβ_1-42_-Cu^2+^, indicating a pronounced enhancement of catechol oxidase-like activity upon membrane binding (**Figure 4B**).

**Figure 4.**
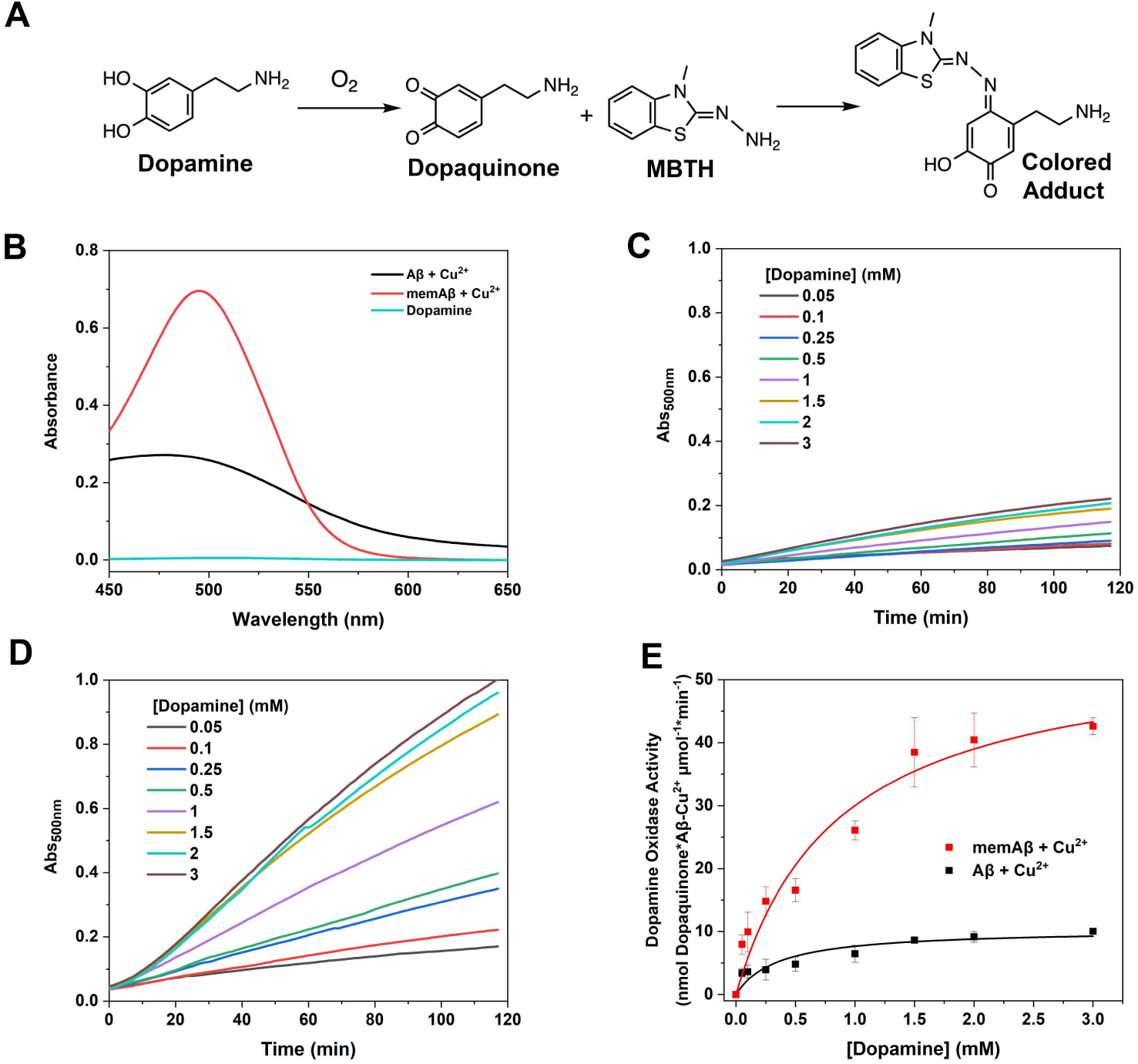
Catechol oxidase activity of soluble and membrane-bound Aβ_1-42_-Cu^2+^ complexes. A) Dopamine oxidation to dopaquinone can be quantified using the spectrophotometric probe MBTH, which forms a colored adduct with dopaquinone (ε_500_ = 32,500 M^−1^ cm^−1^)^22,47,48^. B) Absorption spectrum of the dopaquinone-MBTH adduct recorded after 85 min incubation at room temperature for soluble Aβ_1-42_-Cu^2+^ (10 μM; black), memAβ_1-42_-Cu^2+^ (12 μM; red) in the presence of 2 mM dopamine and 3 mM MBTH (dopamine autoxidation presented in cyan), in 20 mM N-ethylmorpholine pH 7.4, 100 mM NaCl (Aβ_1-42_ samples corrected for dopamine auto-oxidation) C) Representative concentration-dependent kinetic traces at different dopamine concentrations (0.05-3 mM) for soluble Aβ_1-42_-Cu^2+^ (10 μM) (n = 3 ± s.d.) and (D) memAβ_1-42_-Cu^2+^ (10 μM). E) Michaelis-Menten analysis of the dopamine oxidase enzymatic activity for the soluble Aβ_1-42_-Cu^2+^ (black) and memAβ-Cu^2+^ (red), as a function of dopamine concentration (n = 3 ± s.d.).

Dopamine oxidation for memAβ_1-42_-Cu^2+^ and Aβ_1-42_-Cu^2+^ followed an almost linear dependency as a function of time until the reagents started to become limiting (**Figure 4 C-D**). Kinetics of dopamine oxidation as a function of substrate concentration for memAβ_1-42_-Cu^2+^ exhibited a hyperbolic dependency similar to its soluble counterpart (**Figure 4E**). Michaelis-Menten analysis yielded V_max_ values of 55.6 ± 6.8 nmol dopaquinone* Aβ-Cu^2+^ μmol^-1^*min^-1^ for memAβ_1-42_-Cu^2+^, and 10.3 ± 1.2 nmol dopaquinone* Aβ-Cu^2+^ μmol^-1^*min^-1^ with corresponding K_m_ values of 0.8 ± 0.2 mM and 0.3 ± 0.1 mM. These findings demonstrate that membrane insertion significantly amplifies the dopamine oxidase activity of Aβ_1-42_-Cu^2+^, leading to complexes with comparable K_m_ values but V_max_ values approximately five-fold higher than the soluble Aβ_1-42_-Cu^2+^counterpart (**Table 1**).

**Table 1.**
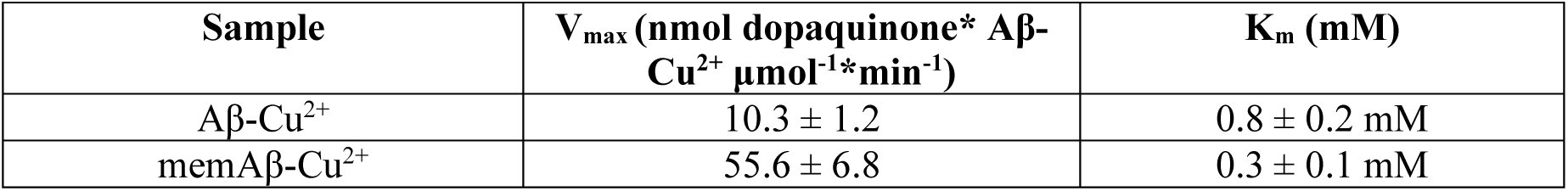
Michaelis-Menten parameters for the enzymatic catechol oxidase activity of soluble and membrane-bound Aβ_1-42_-Cu^2+^ complexes in the presence of 3 mM of MBTH in 20 mM N-ethylmorpholine, 100 mM NaCl, pH 7.4.

| Sample | $V_{\max}$ (nmol dopaquinone* Aβ-Cu$^{2+}$ $\mu\text{mol}^{-1}\cdot\text{min}^{-1}$) | $K_m$ (mM) |
| --- | --- | --- |
| Aβ-Cu$^{2+}$ | $10.3 \pm 1.2$ | $0.8 \pm 0.2$ mM |
| memAβ-Cu$^{2+}$ | $55.6 \pm 6.8$ | $0.3 \pm 0.1$ mM |

Soluble Aβ_1-42_-Cu^2+^ was proposed to act as a Type-3 Cu enzyme.^22^ These enzymes are characterized by possessing a dinuclear Cu center, where three histidine residues coordinate each Cu ion.^49^ Type 3 Cu-enzymes feature an active site capable of binding molecular oxygen and both Cu^2+^ are antiferromagnetically-coupled showing no EPR activity.^49,50^ Catechol oxidases promotes the transformation of catechols to quinones in a cyclic process, where the catechol will coordinate to the active site of the enzyme, leading to the catechol oxidation and reduction of two Cu^2+^ to Cu^+^. Molecular oxygen will bind this reduced dimetallic center leading to the formation of a side-on peroxo bridge with the dimetallic Cu^2+^ center, allowing turnover oxidation of catechols.^22,49,51^ This detrimental interaction could explain some of the pathological hallmarks observed in AD, as the Aβ_1-42_-Cu^2+^ complex would be able to oxidize key neurotransmitters with catechol functional groups such as dopamine, norepinephrine, and epinephrine.^22–24^

We further investigated dopamine oxidation rates across varying dopamine concentrations in the presence of increasing concentrations of H₂O₂. The addition of H₂O₂ resulted in a hyperbolic dependence of dopamine oxidation rates, consistent with a catalytic metal-centered oxidation process, with reaction rates increasing progressively up to approximately 10-20 mM H₂O₂ (**Figure 5**). Analysis of apparent V_max_ changes as a function of H₂O₂ concentration revealed a Michaelis-Menten relationship, yielding V_max,H₂O₂_ of 20.9 ± 4.0 nmol dopaquinone* Aβ-Cu^2+^ μmol^-1^*min^-1^ and K_m,H₂O₂_ of 1.5 ± 2.0 mM for soluble Aβ_1-42_-Cu^2+^, V_max,H₂O₂_ of 112.66 ± 19.8 nmol dopaquinone* Aβ-Cu^2+^ μmol^-1^*min^-1^ and K_m,H₂O₂_ of 0.5 ± 1.3 mM for memAβ_1-42_-Cu^2+^ (**Figure 5**). This hyperbolic dependency suggests that peroxide directly participates in the catalytic mechanism at the metal center when H₂O₂ is supplemented. (For V_max_ values as a function of [H_2_O_2_], see **Supplementary Table S1**) As discussed, it was previously observed for Aβ-Cu^2+^ complexes, when oxidizing THB,^22–24^ that the oxidation could proceed via transient dinuclear Cu^2+^ centers, facilitating two-electron transfers to generate Cu^+^ and ortho-quinone products. The resulting Cu^+^ species could then bind O₂, forming a dinuclear Cu^2+^-peroxo center that oxidizes a second substrate, completing the catalytic cycle in a manner reminiscent of Type 3 Cu centers typical of catechol oxidases. H₂O₂ was shown to bypass part of this cycle, accelerating oxidation turnover. Future studies will be needed to determine whether dopamine oxidation by membrane-bound memAβ_1-42_-Cu^2+^ follows a mononuclear catalytic mechanism or involves transient dinuclear Cu^2+^ centers, akin to soluble Aβ-Cu^2^. Nonetheless, our findings highlight the enhanced oxidative potential of membrane-bound Aβ_1-42_-Cu^2^ compared to its soluble form, underscoring the relevance of these membrane-associated species in the context of copper-mediated neurodegenerative processes. Notably, membrane-bound Aβ_1-42_-Cu^2+^ exhibits catalytic parameters that are lower but approaching the ones of human catechol oxidases such as tyrosinase.^52^

**Figure 5.**
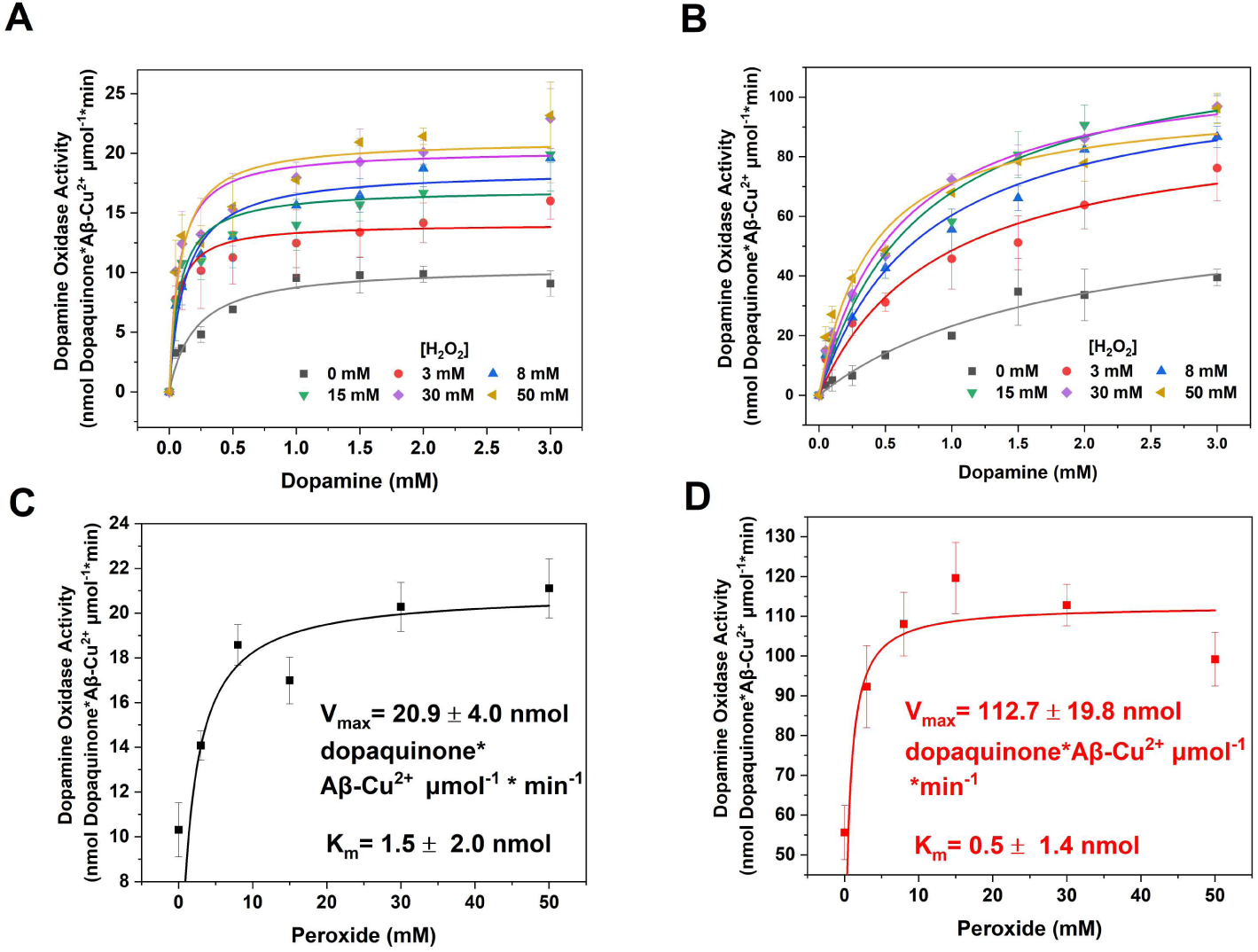
Catechol oxidase activity by Aβ-Cu^2+^ complexes in the presence of H_2_O_2._ Michaelis-Menten kinetic analysis as a function of dopamine concentration (as substrate) at increasing H_2_O_2_, measured in 20 mM N-ethylmorpholine pH 7.4, 100 mM NaCl, and 3 mM MBTH at room temperature: A) soluble Aβ_1-42_-Cu^2+^ (10 μM) and B) memAβ_1-42_-Cu^2+^ (10 μM) (n = 3 ± s.d.). Michaelis-Menten analysis of V_max_ as a function of H_2_O_2_ concentration (as substrate) is presented for C) soluble Aβ_1-42_-Cu^2+^ (12 μM) and D) memAβ_1-42_-Cu^2+^ (12 μM) (n = 3 ± s.d.).

### 3.3. MemAβ-Cu^2+^ redox reactivities: Aβ di-tyrosine crosslinking and hydroxyl radical production

In the presence of biological reducing agents (e.g. ascorbate) and molecular oxygen, Aβ-Cu^2+^ complexes can undergo a Cu^+^/Cu^2+^ redox cycling catalyzing the generation of ROS such as HO_2_^•^ (perhydroxyl radical), O_2_^•−^ (superoxide radical anion), H_2_O_2_ (hydrogen peroxide,) and ^•^OH (hydroxyl radical) via Fenton and Haber-Weiss-type reactions.^28^ These species can have detrimental repercussion against proteins, DNA, and lipids, in particular polyunsaturated fatty acids (PUFAs), that are highly susceptible to peroxidation.^53–55^

The formation of di-tyrosine crosslinks, including in Aβ, has been proposed as a possible biomarker of the oxidative stress of AD.^56,57^ Di-tyrosine has been found in the cerebrospinal fluid (CSF) of AD patients, as well as in AD brains.^57–59^ It has been demonstrated that the presence of Aβ dimers can compromise the process of synaptic plasticity.^58,59^

In Aβ, di-tyrosine cross-links are formed through ortho-ortho coupling of two Tyr10 residues in the presence of oxygen and the reducing agent ascorbate, leading to homodimerization that can serve as a nucleation point for oligomerization and potential fibrillation. This process is thought to be facilitated via copper-mediated electron transfer in Aβ-Cu^2+^ complexes, where oxygen facilitates the abstraction of the phenolic hydrogen, followed by radical tautomerization within the aromatic ring. The resulting tyrosyl radicals subsequently couple to form di-tyrosine cross-links.^56^

To investigate whether Aβ membrane insertion affects or abolishes copper-catalyzed di-tyrosine crosslinking, we examined memAβ_1-42_-Cu^2+^ di-tyrosine formation compared to soluble Aβ_1-42_-Cu^2+^(**Figure 6**). Di-tyrosine adducts exhibit a distinct and characteristic fluorescence emission band centered at 425 nm (λ_ex_ = 325 nm), which can be used to monitor their formation. Incubation of soluble Aβ_1-42_-Cu^2+^ and mem Aβ_1-42_-Cu^2+^ with 3 mM ascorbate for 30 minutes resulted in a marked increase in fluorescence emission characteristic of di-tyrosine formation (**Figure 6 A-C**). In contrast, negligible fluorescence emission was observed in control samples lacking Cu^2+^ when compared to soluble Aβ_1-42_-Cu^2+^ and memAβ_1-42_-Cu^2+^. Kinetic analysis at 425 nm further confirmed a progressive increase in the di-tyrosine emission signal, which was absent in control samples, supporting a metal-dependent cross-linking mechanism (**Figure 6D**). Thus, the binding to membrane does not prevent di-tyrosine crosslinks generation in memAβ_1-42_-Cu^2+^, potentially facilitating this detrimental reaction when the peptide is embedded into the 2-dimensional environment of the lipid bilayer.

**Figure 6.**
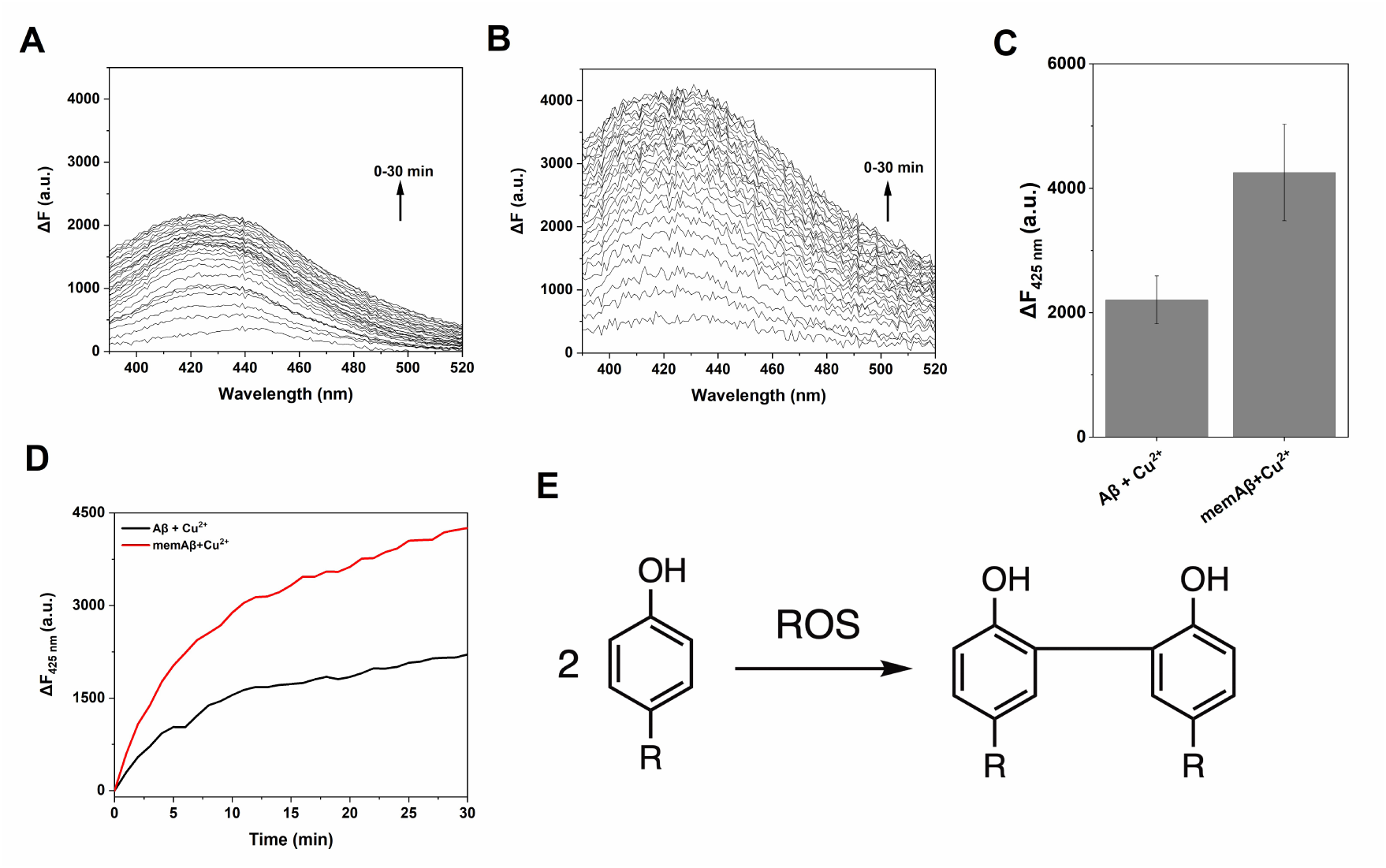
Di-tyrosine crosslink formation catalyzed by Aβ-Cu^2+^ complexes. A) Fluorescence emission spectrum (λ_ex_ = 325 nm) reporting di-tyrosine formation (minus the contribution of the equivalent samples in absence of Cu^2+^) for (A) soluble Aβ_1-42_-Cu^2+^ (10 μM; black) and (B) memAβ_1-42_-Cu^2+^ (10 μM; red), recorded for 30 min at 37 °C in 20 mM phosphate buffer, pH 7.4, and 3 mM ascorbate. C) Quantification of di-tyrosine formed by Aβ-Cu^2+^ complexes obtained from the change in fluorescence at 425 nm (ΔF_425nm_). Values obtained in samples without Cu^2+^ for both soluble and membrane-bound species were used as background, and their contribution subtracted (n = 3 ± s.d.). C) Kinetic traces following the formation of di-tyrosine crosslinks for Aβ-Cu^2+^ complexes (soluble, black; membrane-bound, red). E) Scheme showing the ortho-ortho cross coupling in the benzylic ring of two tyrosine residues upon their interaction with ROS.

The results obtained substantiate that memAβ_1-42_-Cu^2^ possess remarkable catalytic redox activities including radical-centered chemistry. Copper redox-cycling via Fenton and Haber Weiss-like reactions with molecular oxygen and biological reducing agents represent a dramatic potentiating factor in promoting Aβ toxicity in the AD brain. It is worth highlighting that, under physiological conditions, in the central nervous ascorbate is present in high concentrations, up to 10 mM in neurons.^14^ To additionally investigate this redox chemistry for memAβ_1-42_-Cu^2+^, we determined catalytic ROS production by quantifying hydroxyl radical generation by fluorescence spectroscopy using the reporter probe coumarin-3-carboxylic acid (3-CCA). 3-CCA acts as a hydroxyl radical scavenger, producing the fluorescent compound 7-hydroxycoumarin-3-carboxylic acid (7-OH-3-CCA; λ_ex_=395 nm, λ_em_=450 nm) upon hydroxylation.^60^ Due to its non-fluorescent nature and high hydroxylation rate constant, detecting 7-OH-3-CCA enables real-time monitoring of hydroxyl radical generation by Aβ_1-42_-Cu^2+^ complexes (**Figure 7A**). We monitored hydroxyl radical formation kinetics in samples containing soluble Aβ_1-42_-Cu^2+^ or memAβ_1-42_-Cu^2+^ (10 μM) with ascorbate (1.2 mM) and compared their catalytic efficiency to those of the peptides in samples without Cu^2+^ (**Figure 7B-C**). Fluorescence analysis showed that both Aβ_1-42_-Cu^2+^ complexes effectively catalyzed ascorbate-driven hydroxyl radical production at significant rates, also considering that lipid molecules in the reaction of memAβ_1-42_-Cu^2+^ can partially contribute to radical quenching. Overall, all these findings demonstrate that membrane-bound Aβ_1-42_-Cu^2+^ complexes exhibit significant oxidative activity when compared to their soluble counterparts, highlighting the crucial role of these membrane-associated species when investigating copper-mediated neurodegenerative mechanisms in AD.

**Figure 7.**
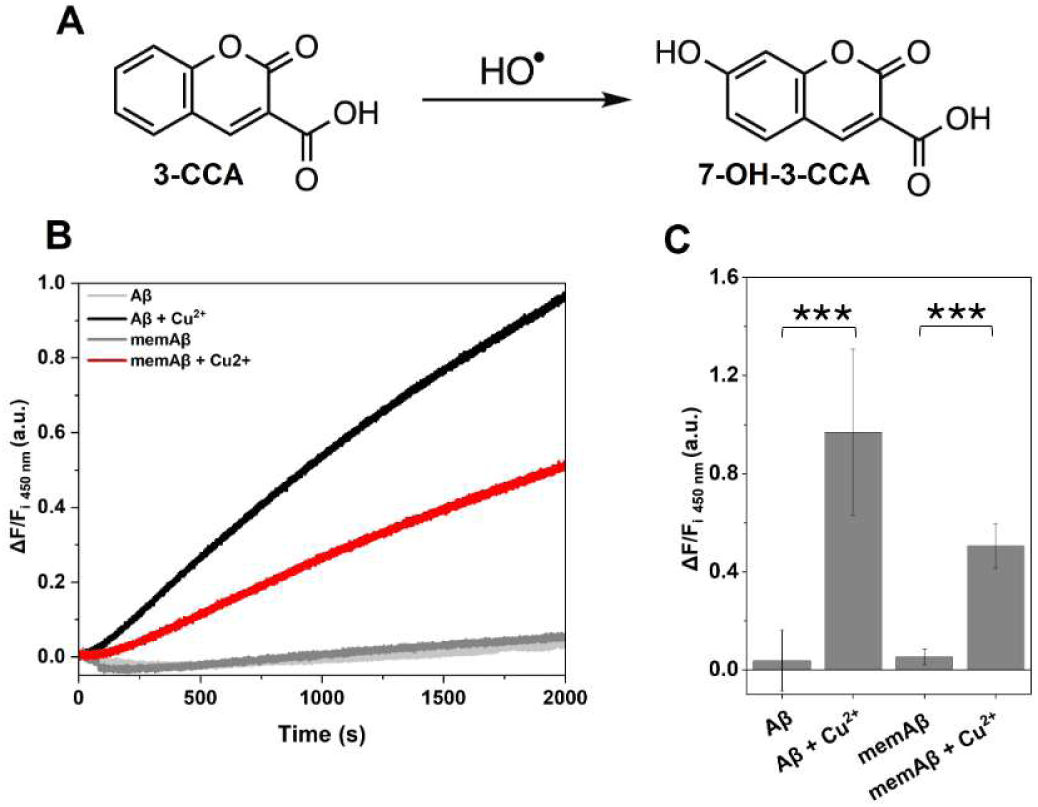
Ascorbate-driven hydroxyl radical production by Aβ-Cu^2+^ complexes. A) Hydroxyl radical reaction with the fluorescent probe 3-CCA yielding the formation of the fluorescent 7-OH-3-CCA product. B) Hydroxyl radical generation kinetics for soluble Aβ_1-42_-Cu^2+^ (10 μM, black) (n = 7 ± s.d.) and memAβ_1-42_-Cu^2+^ (10 μM, red) (n = 4 ± s.d.) in the presence of 1.2 mM ascorbate, in 20 mM phosphate buffer pH 7.4 at 37 °C, followed by 7-OH-3-CCA fluorescence emission. C) Relative quantification of hydroxyl radicals generated by Aβ-Cu^2+^ complexes (30 min) obtained from the fluorescence change over the initial fluorescence at 450 nm (ΔF/F_i 450nm_). For statistical analysis, a student’s t test was performed (95% confidence): P>0.05 (ns), P≤0.05 (*), P≤10^-2^ (**), P≤10^-3^ (***), P≤10^-4^ (****).

### 3.4 MemAβ-Cu^2+^ complexes catalyze lipid peroxidation and disrupt membrane integrity

In the CNS of AD patients extensive lipid peroxidation is observed as a pathophysiological hallmark, significantly contributing to the neurodegenerative processes.^61^ The role of membrane-bound Aβ species in promoting this processes has not been investigated in detail. Cells possess protective mechanisms to prevent lipid peroxidation damage via the action of antioxidant proteins and small molecules, but such mechanisms can be overcome when the concentration of free radicals is high, leading to cell death and/or apoptosis.^62^

The process of lipid peroxidation of a PUFA occurs in three key steps: 1) initiation, 2) propagation, and 3) termination. The reactions starts when an allylic hydrogen is abstracted by one of the radical species leading to the formation of a carbon-centered lipid radical that can undergo rearrangement.^62–65^ Molecular oxygen can then rapidly react with a lipid radical forming a highly reactive^66^ lipid peroxyl radical, as well as lipid hydroperoxydes.^65^ After two cyclization processes, an endoperoxide is generated that will lead to the formation of malondialdehyde (MDA; **Figure 8**), a central lipid peroxidation end-products.^65^ MDA can form adducts with several biomolecules, including the DNA and RNA nucleotide bases, as well as proteins.^67^ This species is considered the most mutagenic product generated from PUFA lipid peroxidation.^62,67^ MDA has been extensively used as a marker of lipid peroxidation.^67^ Neurons are particularly susceptible to toxicity from lipid peroxidation end-products, which could account for some of the hallmarks of AD.^61^

**Figure 8.**
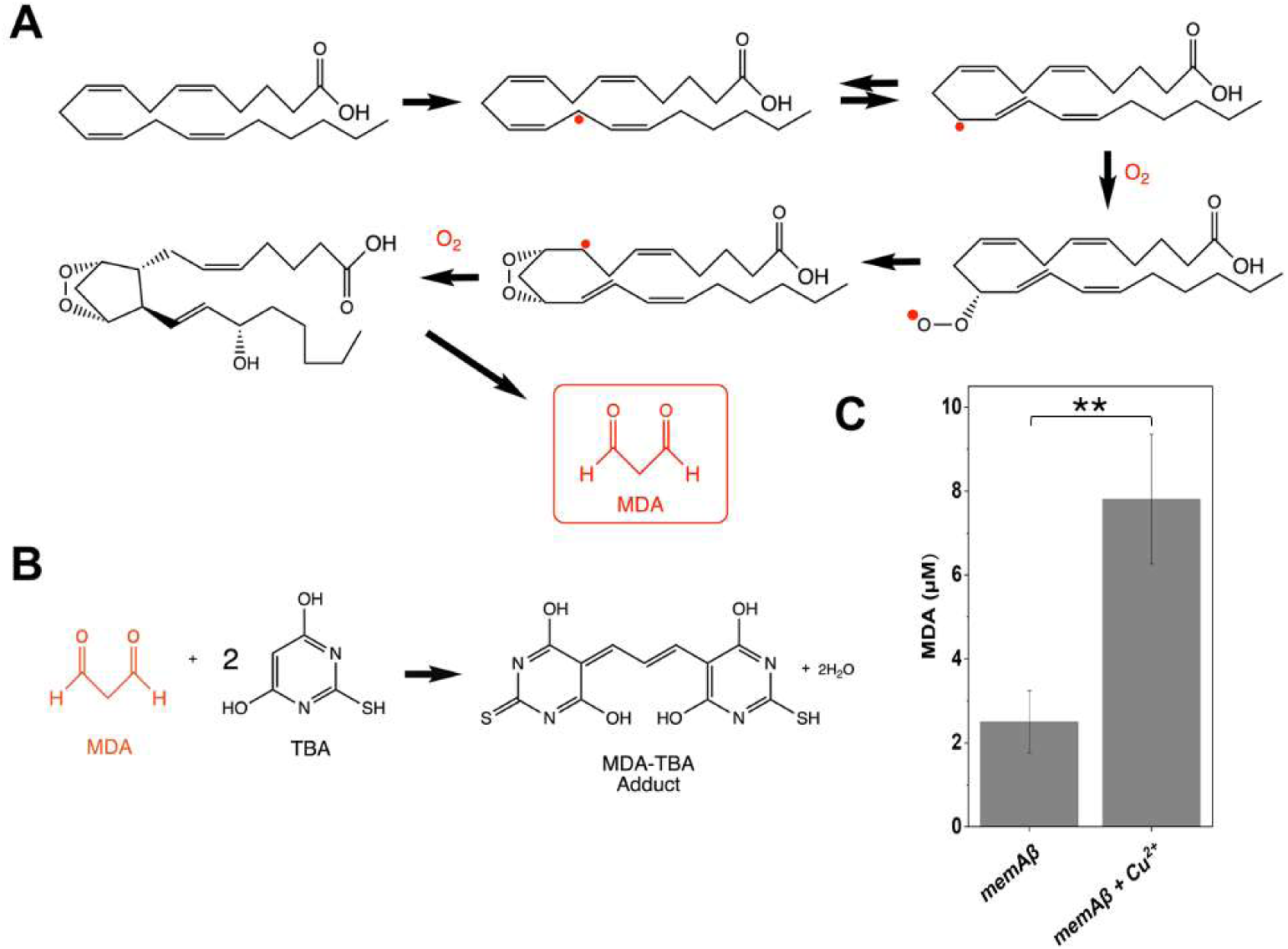
Generation of malondialdehyde (MDA) from PUFA peroxidation. A) Reaction scheme presenting arachidonic acid (AA) peroxidation producing malondialdehyde (MDA). First, a radical abstracts a hydrogen from the aliphatic tail double bound forming a lipid radical, that can undergo rearrangement. Reaction with oxygen leads to a lipid peroxyl radical promoting a cyclization process. Further reaction with another oxygen molecule generates the MDA end-product via retro-Diels-Alder reaction.^69^ B) MDA reaction with 2-thiobarbituric acid (TBA) yielding a colored product which can be quantified at λ_abs_ = 532 nm.^70^ C) MDA quantification produced by memAβ (12 μM) using SUVs containing 1% (w/w) AA (AALip), in the presence or absence of Cu^2+^, recorded in 20 mM phosphate buffer pH 7.4. Ascorbate (1mM) was added every hour for 6 hours at 37 °C, under agitation. After 10 hours, 4 more additions of ascorbate were performed in a 4-hour window. The samples were centrifuged, and the absorption at 532 nm in supernatant was measured. The memAβ_1-42_-Cu²⁺ species lead to significantly higher lipid peroxidation (n = 4 ± s.d.). A sample exclusively consisting of SUVs (AALip) was used as a reference to correct absorbance values. For statistical analysis, a student’s t test was performed (95% confidence): P>0.05 (ns), P≤0.05 (*), P≤10^-2^ (**), P≤10^-3^ (***), P≤10^-4^ (****).

As we established that memAβ_1-42_-Cu²⁺ feature a remarkable capability to perform redox catalysis and efficiently generate ROS, we hypothesized that oxidative radical species could efficiently react with the unsaturated bonds of PUFAs within the two-dimensional lipid bilayer (where memAβ_1-42_-Cu²⁺ can laterally diffuse). PUFAs, being key structural components of CNS cellular membranes, are particularly susceptible to lipid peroxidation. To investigate this reactivity, we generated SUVs in which 1% arachidonic acid (AA) was doped into the lipid bilayer (POPC:POPG:AA, 69.3:29.7:1; w/w/w). AA was selected due to its critical role in cellular signaling and neuronal integrity maintenance.^68^ The association and membrane insertion of Aβ_1-42_ into SUVs featuring this modified lipid formulation (AALip) was confirmed by size-exclusion chromatography (SEC) and SDS PAGE analysis upon silver staining (**Supplementary Figures S1-2**).

To evaluate ROS-induced lipid peroxidation, we supplemented ascorbate to memAβ_1-42_-Cu²⁺ SUVs containing AA and quantified MDA formation over time, as biomarker of oxidative lipid peroxidation. MDA was detected using a thiobarbituric acid (TBA) assay, which produces a chromogenic adduct that absorbs in the visible spectral range, allowing to quantify oxidative stress-related lipid peroxidation (**Figure 8**).^64–67^ Comparative analysis between memAβ_1-42_ samples with and without Cu²⁺ revealed a significantly increase in MDA production and lipid peroxidation upon formation of memAβ-Cu²⁺ complexes (**Figure 8**). This work demonstrates that memAβ_1-42_-Cu²⁺ can act as an efficient catalyst for detrimental oxidation of biological membranes, potentially contributing to lipid bilayer destabilization and disruption of the membrane structural integrity, possibly leading to membrane leakage.

As memAβ_1-42_-Cu²⁺ catalyze lipid peroxidation, we further examined whether this reactivity could promote membrane permeabilization and leakage in SUVs. To monitor membrane stability, we encapsulated the fluorescent probe 5(6)-carboxyfluorescein within the AALip SUV lumen. Samples were sonicated, and vesicle size distribution determined using DLS, confirm satisfactory size and polydispersity indexes (**Figure S3**) after encapsulation. Unencapsulated fluorescent probe was removed by centrifugation and the vesicles were incubated with Aβ_1-42_, followed by addition of Cu²⁺ to generate memAβ_1-42_-Cu²⁺, and the reaction initiated by the addition of ascorbate. After ultracentrifugation to pellet SUVs, fluorescence 5(6)-carboxyfluorescein emission spectra (F_i_) were recorded from 500 to 650 nm (λ_ex_ = 450 nm) in the supernatant fraction to quantify the leaked probe. In parallel the vesicle pellet was resuspended, and complete membrane disruption was induced by the addition of detergent, allowing complete release of the unleaked probe, which allowed for quantification by fluorescence emission (F_f_). To determine the relative amount of leaked probe in each sample, the area under the 5(6)-carboxyfluorescein fluorescence emission band was integrated, and the relative fluorescence was determined upon calculating the ratio of F_i_ to the total fluorescence (F_i_ + F_f_). This analysis demonstrated that the generation of memAβ_1-42_-Cu²⁺ complexes resulted in a significant increase in probe leakage, strongly indicating that membrane Aβ_1-42_-Cu²⁺ insertion triggers membrane destabilization, disrupting structural integrity and leading to lipid bilayer leakage (**Figure 9**).

**Figure 9.**
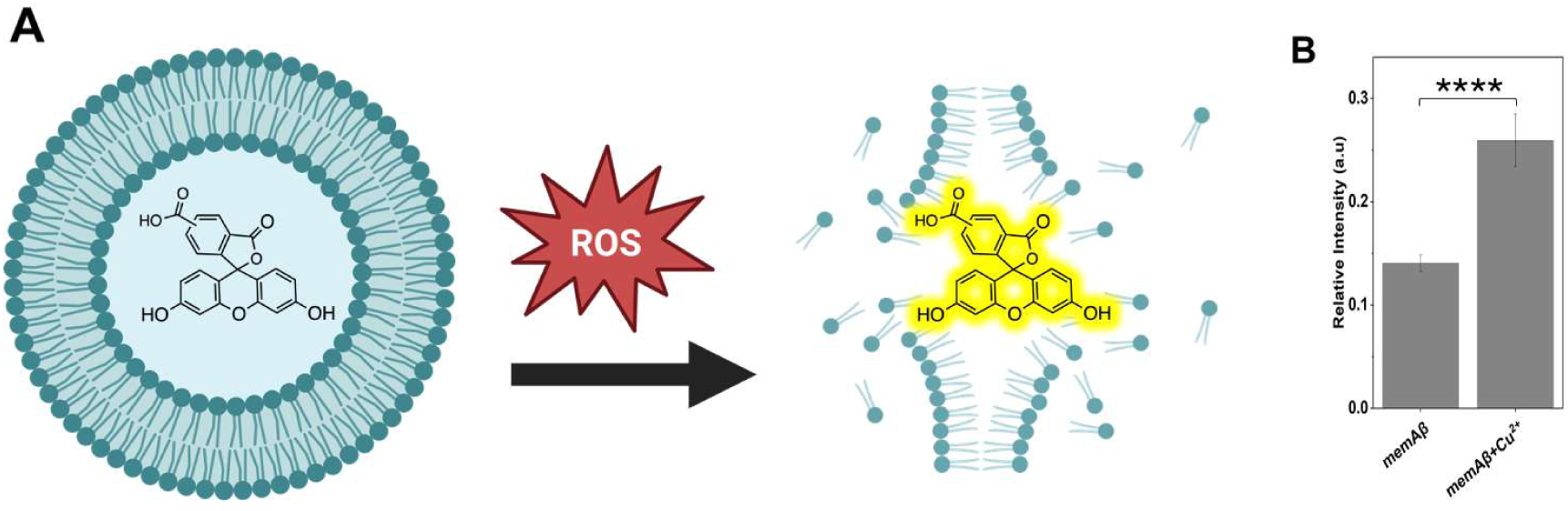
Membrane leakage promoted by memAβ-Cu^2+^. A) Cartoon showing the fluorescent probe (5(6)-carboxyfluorescein) encapsulated inside the lumen of SUVs containing 1% (w/w) arachidonic acid. ROS generated by memAβ_1-42_-Cu^2+^ can lead to membrane structure destabilization and leakage. B) memAβ-Cu^2+^ promotes the generation of ROS compromising the integrity of the artificial lipid bilayer. The area under the curve of the fluorescence spectrum (λ_ex_= 450 nm; λ_em_ = 500-650 nm) was recorded for the supernatant and the SUV pellet upon ultracentrifugation (after the addition of Triton X-100 to solubilize the SUV pellet). The relative fluorescence intensity was calculated as reported in The Methods section. The fluorescence contribution of samples containing exclusively lipids (in absence of ascorbate) was subtracted from all the samples (n = 9 ± s.d.). For statistical analysis, a student’s t test was performed (95% confidence): P>0.05 (ns), P≤0.05 (*), P≤10^-2^ (**), P≤10^-3^ (***), P≤10^-4^ (****).

Overall, this established a key correlation between Aβ_1-42_-Cu²⁺ redox catalytic properties upon lipid bilayer insertion and detrimental chemistry occurring in the membranes, establishing that memAβ_1-42_-Cu²⁺ can efficiently trigger and potentiate neuronal toxicity by altering normal transmembrane impermeability and the associated transmembrane potential.

### 3.5 Zn_7_MT-3 efficiently quenches all redox reactivities of memAβ-Cu^2+^, prevents lipid peroxidation, and promotes membrane stability

Copper coordination by amyloidogenic proteins, including Aβ, has been demonstrated to be a key contributing factor to neurodegenerative processes due to their well-documented redox activities. Given the crucial function of metallothionein-3 (MT-3) in regulating copper homeostasis and preventing aberrant copper-protein interactions in the central nervous system,^38^ as demonstrated for soluble Aβ-Cu^2+^ species,^26^ we explored in this study the reactivity between Zn₇MT-3 and memAβ_1-42_-Cu²⁺. Zn₇MT-3, which is present both intra- and extracellularly in the CNS, was identified as a potential suppressor of Aβ-Cu^2+^ toxicity in AD. This Zn₇MT-3 activity stems from its ability scavenge Cu^2+^, reducing it to Cu^+^ and sequestering it in a redox-inert Cu^+^_4_-thiolate cluster, while forming intramolecular disulfide bonds, thereby preventing ROS generation.^25^ This detoxifying mechanism has been observed in the case of soluble Aβ-Cu^2+^,^26^ as well as other proteins involved in neurodegenerative disorders such as α-Syn (PD).^27,28^

Given that Aβ-Cu^2+^ complexes can exist in both soluble and membrane-bound states and considering that membrane insertion significantly amplifies the dopamine oxidase activity of Aβ_1-42_-Cu^2+^ relative to its soluble counterpart, we aimed to determine whether soluble Zn₇MT-3 can also target these membrane-bound complexes and neutralize their redox toxicity. As MT-3 is a soluble metalloprotein, its observed reactivity towards soluble Aβ_1-42_-Cu^2+^ does not provide direct evidence for whether this interaction also occurs when Aβ-Cu^2+^ is embedded within lipid bilayers.

To assess the impact of Zn₇MT-3 on the dopamine oxidase activity of membrane-bound Aβ_1-42_-Cu^2+^ and compare it to soluble Aβ_1-42_-Cu^2+^, we incubated Aβ_1-42_-Cu^2+^complexes with Zn₇MT-3 (0.25 eq.) at 25 °C for 1 hour and monitored dopamine ortho-quinone formation using MBTH in the presence of 1 mM dopamine. By tracking the absorbance kinetics of the MBTH-quinone adduct at 450–650 nm over an 85-minute period, we observed that Zn₇MT-3 completely abolished the dopamine oxidase activity not only of soluble Aβ_1-42_-Cu^2+^ but also memAβ_1-42_-Cu^2+^, restoring oxidation levels to the baseline of spontaneous dopamine auto-oxidation (**Figure 10**).

**Figure 10.**
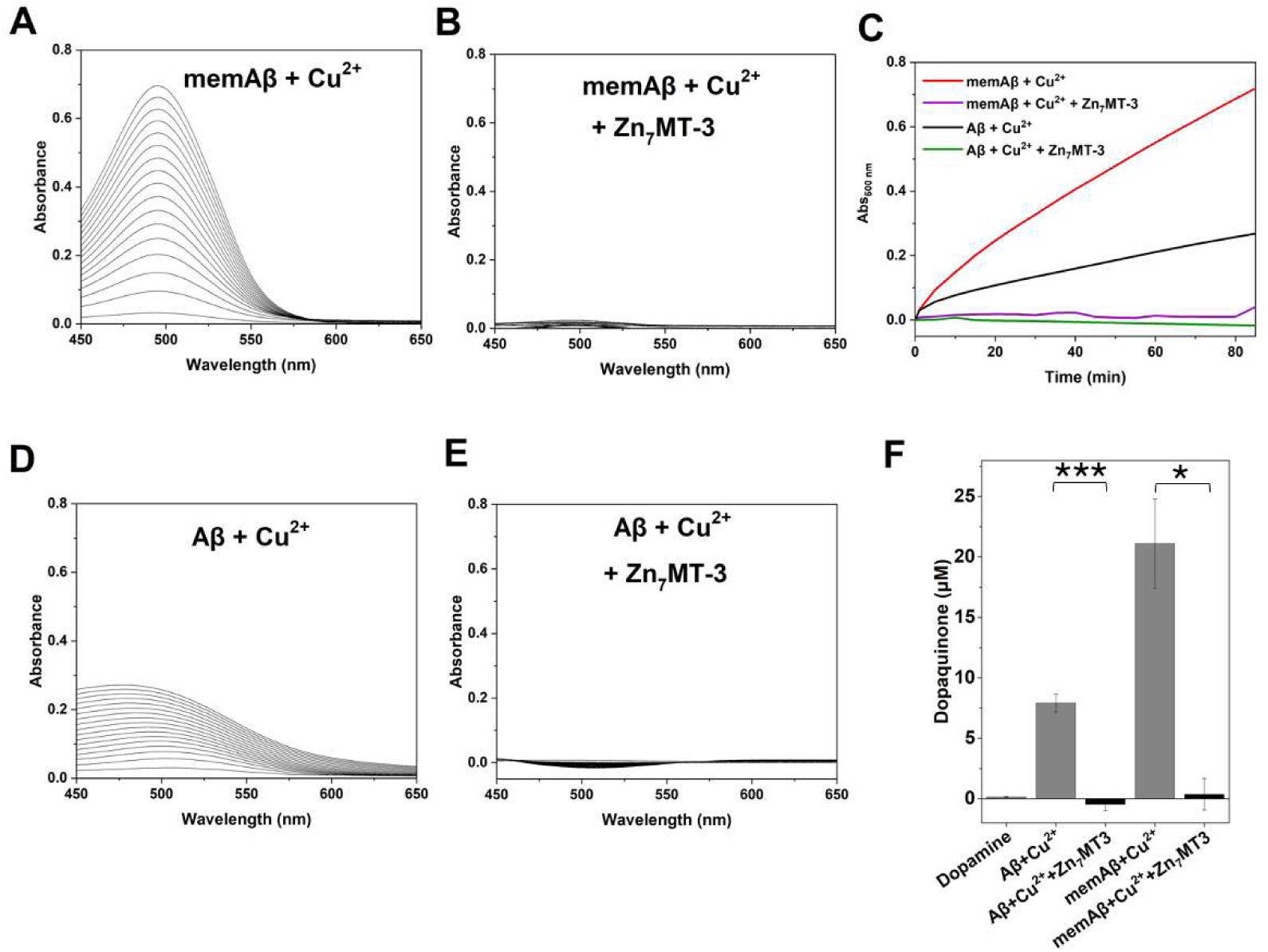
Aβ-Cu^2+^ dopamine oxidase activity upon reaction with Zn_7_MT-3. The MBTH-quinone adduct absorbance spectrum was recorded every 5 min for 85 min for (A) memAβ-Cu^2+^ (10 μM), (B) memAβ-Cu^2+^ plus Zn_7_MT-3 (10 μM: 2.5 μM), (D) soluble Aβ_1-42_-Cu^2+^ (10 μM), and (E) memAβ-Cu^2+^ plus Zn_7_MT-3 (10 μM: 2.5 μM) (n = 3 ± s.d.). (C) Kinetic traces reporting the oxidation of dopamine to dopaquinone in the presence of MBTH: memAβ-Cu^2^ (red), memAβ-Cu^2+^ upon reaction with Zn_7_MT-3 (purple), soluble Aβ-Cu^2+^ (black), and soluble Aβ-Cu^2+^ upon reaction with Zn_7_MT-3 (green). (F) Dopaquinone quantification establishing the ability of Zn_7_MT3 to quench the catechol oxidase activity of soluble and membrane-bound Aβ-Cu^2^ complexes (n = 3 ± s.d.). For statistical analysis, a student’s t test was performed (95% confidence): P>0.05 (ns), P≤0.05 (*), P≤10^-2^ (**), P≤10^-3^ (***), P≤10^-4^ (****).

To elucidate the molecular mechanism underlying the observed catechol oxidase activity quenching effect, we examined whether Zn_7_MT-3 facilitates Cu^2+^ removal from Aβ_1-42_-Cu^2+^ and whether this process involves a redox reaction. According to Pearson’s Hard and Soft Acids and Bases (HSAB) theory, Cu^2+^ preferentially coordinates with “ hard” N/O ligands, whereas Cu^+^ exhibits higher affinity for “soft” thiolate ligands, such as those present in MTs Therefore, we assessed whether Zn_7_MT-3 can reduce Cu^2+^ to Cu^+^ and sequester it within its N-terminal β-domain, forming the Cu^+^_4_Zn^2+^_4_MT-3 complex, as previously demonstrated for soluble Aβ-Cu^2+^, α-Syn-Cu^2+^, and PrP-Cu^2+^ complexes.^26–28,37^

At 77 K, Cu^+^_4_Zn^2+^_4_MT-3 exhibits a characteristic luminescence emission spectrum (λ_ex_ = 320 nm) with two distinct emissive bands centered at 425 nm and 575 nm, with corresponding lifetimes (τ) of approx. 40 and 120-140 μs, respectively.^25^ These spectral features indicate the formation of a Cu^+^_4_-thiolate cluster within the MT-3 β-domain, generated via the transient formation of disulfide radical anion intermediates.^36^ The presence of two emission bands is associated with short Cu···Cu internuclear distances (<2.8 Å), where the lower-energy band at 575 nm arises from a triplet charge transfer (CT) transition, and the higher-energy band at 425 nm corresponds to a cluster-centered (CC) transition.^26^ The existence of this high-energy band suggests significant metal-metal interactions, enabling d^10^-d^10^ orbital overlap and metal-metal bonding, which confer remarkable stability against molecular oxygen—an uncommon property for Cu^+^-thiolate clusters—crucial for redox silencing.^35^ To investigate Cu^2+^ reduction and sequestration, we incubated soluble Aβ_1-42_-Cu^2+^ and memAβ_1-42_-Cu^2+^ (10 μM) with Zn_7_MT-3 (2.5 μM) for 1 h before separating membrane-Aβ_1-42_ vesicles (pellet, PT) by ultracentrifugation, and rapidly freezing the MT-3-containing supernatant (SN) samples (as confirmed by SDS-PAGE analysis, *vide infra*) in liquid nitrogen to record their emission spectra at 77 K (λ_ex_ = 320 nm). The resulting spectra exhibited two emission bands with identical peak positions and lifetimes as those of Cu^+^_4_Zn^2+^_4_MT-3, indicating efficient Cu^2+^ removal from memAβ_1-42_-Cu^2+^, reduction to Cu^+^, and incorporation into a redox-inert Cu^+^_4_-thiolate within MT-3 (**Figure 11 and Supplementary Table S2**).

**Figure 11.**
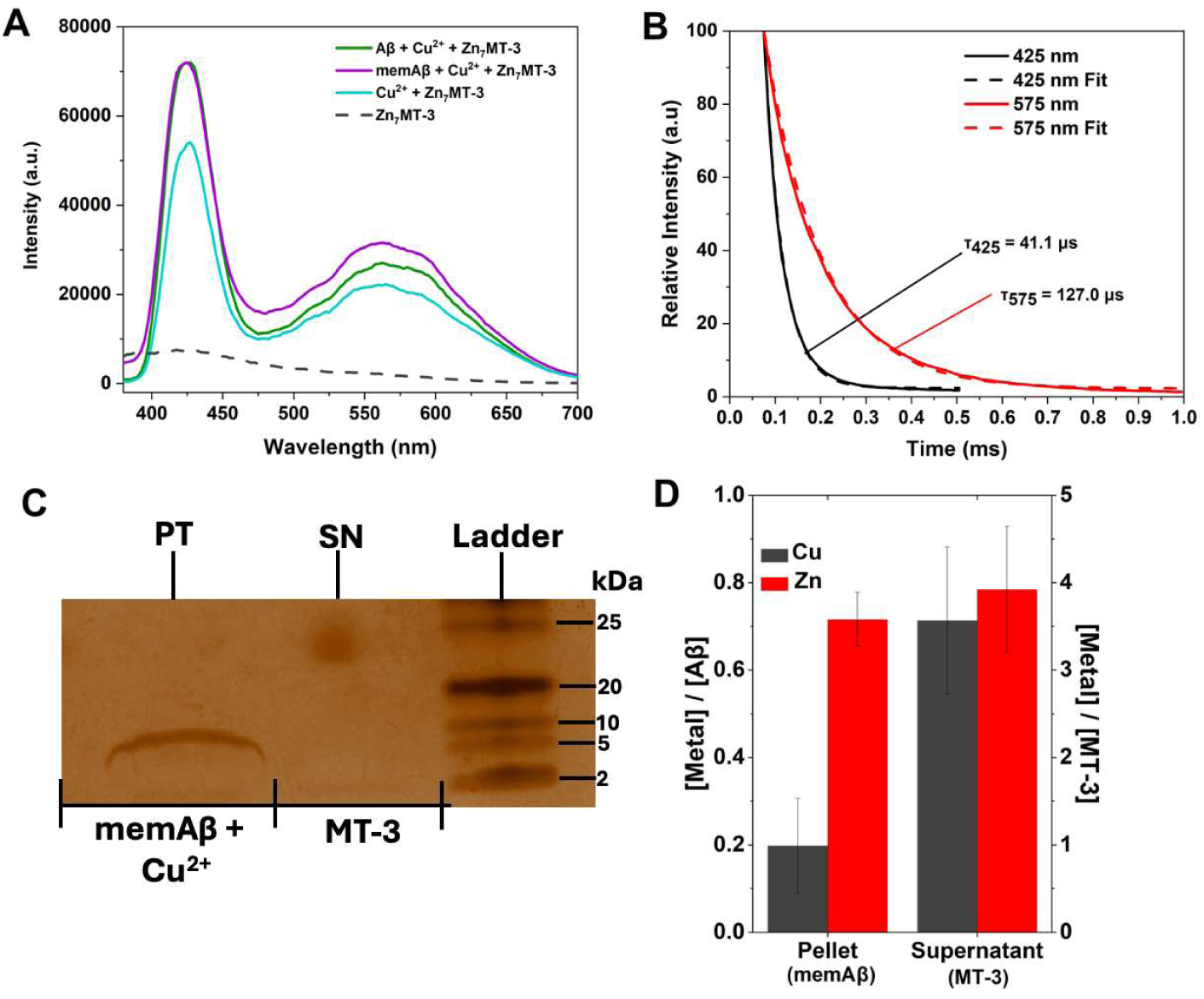
Metal swap reaction between memAβ-Cu^2+^ and Zn_7_MT-3. A) 77 K emission phosphorescence spectrum (λ_ex_ = 320 nm) establishing the formation of the Cu^+^_4_-thiolate cluster in MT-3 upon reaction of Zn_7_MT-3 (2.5 μM) with memAβ_1-42_-Cu^2+^ (10 μM: purple) in 20 mM N-ethylmorpholine pH 7.4, 100 mM NaCl, using the reactions of soluble Aβ_1-42_-Cu^2+^ (10 μM; green) and Cu^2+^ (10 μM: blue) with Zn_7_MT-3 (2.5 μM) as controls. The high energy band at 425 nm and the low energy band at 575 nm are diagnostic of the formation of the Cu^+^_4_-thiolate cluster in Cu^+^_4_Zn^2+^_4_MT-3. B) Exponential decay fitting for the 425 and 575 nm emission bands of Cu^+^_4_Zn^2+^_4_MT-3 generated upon reaction between memAβ-Cu^2+^ with Zn_7_MT-3. The supernatant sample containing Cu^+^_4_Zn^2+^_4_MT-3 was obtained upon ultracentrifugation to remove SUVs. The obtained lifetimes are diagnostic for the presence of the Cu^+^_4_-thiolate cluster in MT-3. C) SDS-PAGE and silver staining demonstrating the presence of memAβ in the SUV pellet and MT-3 as a smear in the supernatant upon the metal swap reaction. D) ICP-MS analysis quantifying Cu/Zn content in MT-3 (SN) in memAβ (PT). The released Zn appears to bind to memAβ_1-42_.

To quantitatively validate this metal scavenging process, we separated via ultracentrifugation membrane-Aβ_1-42_-Cu^2+^ vesicles (PT) from MT-3 (SN) and analyzed metal content by ICP-MS. Consistent with the luminescence data, ICP-MS analysis revealed that >70% of Cu was recovered in supernatant, confirming its sequestration by MT-3. Furthermore, evidence of a partial metal exchange reaction between memAβ_1-42_-Cu^2+^ SUVs and Zn_7_MT-3 was obtained by assessing Zn distribution in the sample. The detection of Zn in the PT post-reaction indicated that Zn^2+^ was released from MT-3 upon Cu^+^ coordination and subsequently bound to memAβ_1-42_ SUVs, further suggesting a metal swap reaction mechanism.

To further validate that Zn_7_MT-3 efficiently reduces and sequesters Cu^2+^, thereby generating the redox-inactive Cu^+^_4_-thiolate complexes and abolishing Aβ_1-42_-Cu^2+^ redox activity, we investigated the impact of these metal swap reactions on ROS production and di-tyrosine cross-linking. Specifically, we assessed di-tyrosine end-product as a marker of oxidative Aβ_1-42_ modifications and monitored hydroxyl radical production using 3-CCA.

To evaluate the effect of Zn_7_MT-3 on di-tyrosine formation, cross-linking in the presence of ascorbate (3 mM) was analyzed and quantified by recording the time-dependent fluorescent emission of di-tyrosine adducts at 425nm. The results revealed a substantial reduction in di-tyrosine formation for both soluble and membrane-bound Aβ_1-42_-Cu^2+^ (**Figure 12 A-B**), further corroborating the abolition of Aβ_1-42_-Cu^2+^ oxidative reactivity.

**Figure 12.**
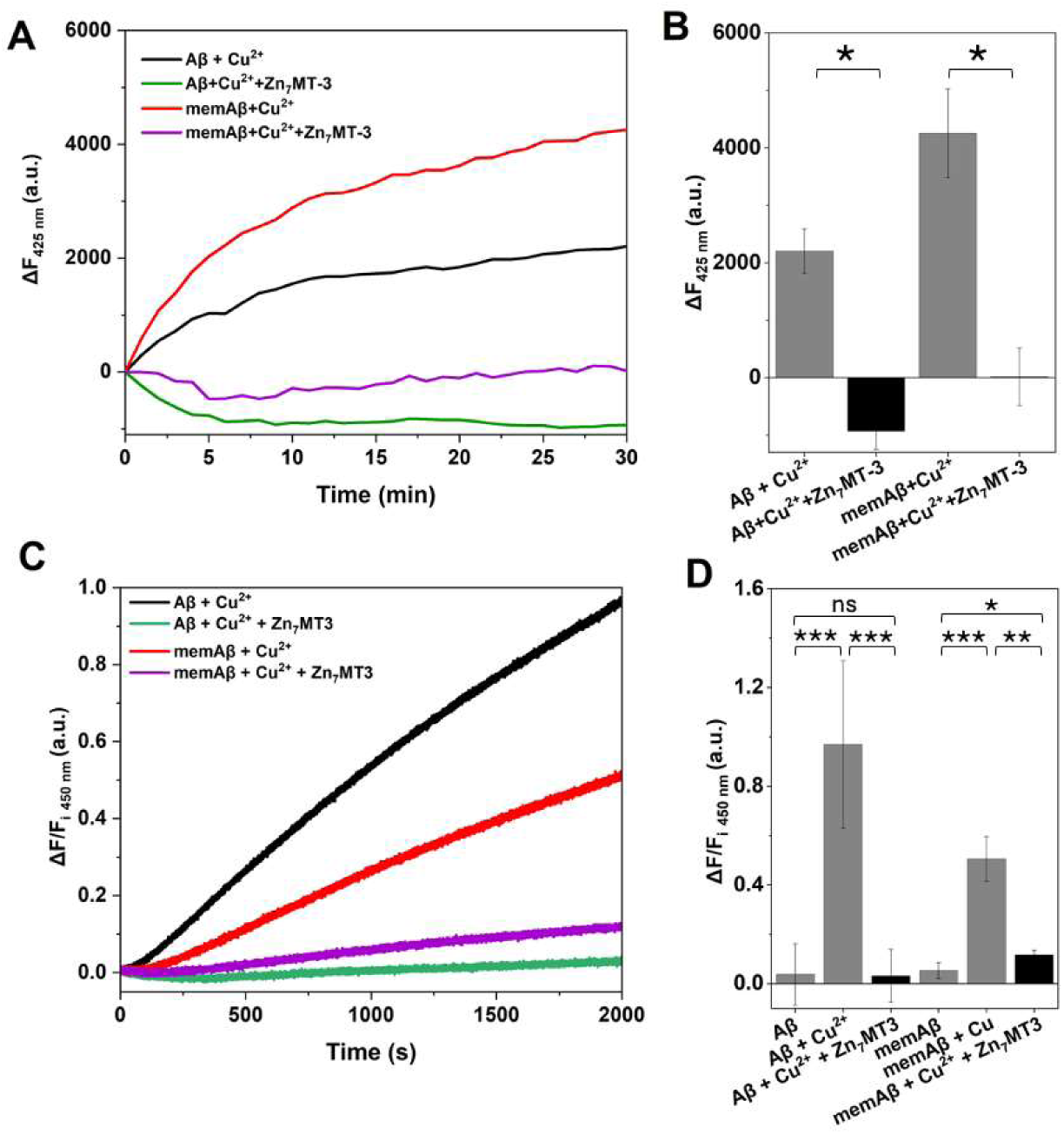
Zn_7_MT-3 quenching of ROS production for all Aβ-Cu^2+^ complexes. A) Di-tyrosine crosslinking reaction kinetics followed by fluorescence emission for memAβ_1-42_-Cu^2+^ (10 μM; red) and soluble Aβ_1-42_-Cu^2+^ (10 μM; black) in the presence of 3 mM ascorbate in 20 mM phosphate buffer, pH 7.4. Upon reaction with Zn_7_MT-3 (2.5 μM) the production of di-tyrosine was completed silenced: memAβ_1-42_-Cu^2+^ plus Zn_7_MT-3 (purple), and soluble Aβ_1-42_-Cu^2+^ plus Zn_7_MT-3 (green). B) Relative di-tyrosine species quantification upon correcting the absorbance of the dopaquinone-MBTH adduct with values obtained in samples without Cu^2+^ and MT-3 (n = 3 ± s.d.). C) Hydroxyl radical production kinetics, followed by fluorescence upon generation of the 7-OH-3-CCA product: Aβ_1-42_-Cu^2+^ (black) (n = 7 ± s.d.), memAβ_1-42_-Cu^2+^ (red), and upon reaction with Zn_7_MT-3 (green and purple, respectively). D) The production of hydroxyl radical species is significantly reduced when Aβ-Cu^2+^ complexes undergo a metal swap reaction with Zn_7_MT-3 (n = 4 ± s.d.). For statistical analysis, a student’s t test was performed (95% confidence): P>0.05 (ns), P≤0.05 (*), P≤10^-2^ (**), P≤10^-3^ (***), P≤10^-4^ (****).

In parallel, to evaluate ROS generation, soluble and membrane-bound Aβ_1-42_-Cu^2+^ complexes were incubated with Zn_7_MT-3 (0.25 eq.), followed by the addition of ascorbate (1.2 mM), and hydroxyl radical production was tracked by monitoring the fluorescence emission of 7-OH-3-CCA (λ_ex_=395 nm, λ_em_=450 nm). Notably, also in this case no significant increase in 7-OH-3-CCA fluorescence was observed (**Figure 12C-D**), indicating complete suppression of hydroxyl radical generation.

Our previous assays demonstrated that memAβ-Cu^2+^ can efficiently and detrimentally catalyze a copper and ROS dependent lipid peroxidation which in turn dramatically affects the integrity of the membrane lipid bilayer. In light of the determined ability of Zn_7_MT-3 to scavenge Cu^2+^ from memAβ-Cu^2+^ and silence it in a redox inert Cu^+^-thiolate cluster, we determined its effect in protecting SUVs containing PUFAs from radical dependent MDA production. Upon analysis of the samples co-incubated with Zn_7_MT-3 a complete quenching of lipid peroxidation was observed (**Figure 13A**), consistent with the described metal swap reaction between Zn_7_MT-3 and memAβ-Cu^2+^ SUVs. Accordingly, membrane leakage assay confirmed the protective role played by MT-3 in maintaining the membrane structural integrity and reducing memAβ-Cu^2+^-catalyzed peroxidation-dependent leakage (**Figure 13B**).

**Figure 13.**
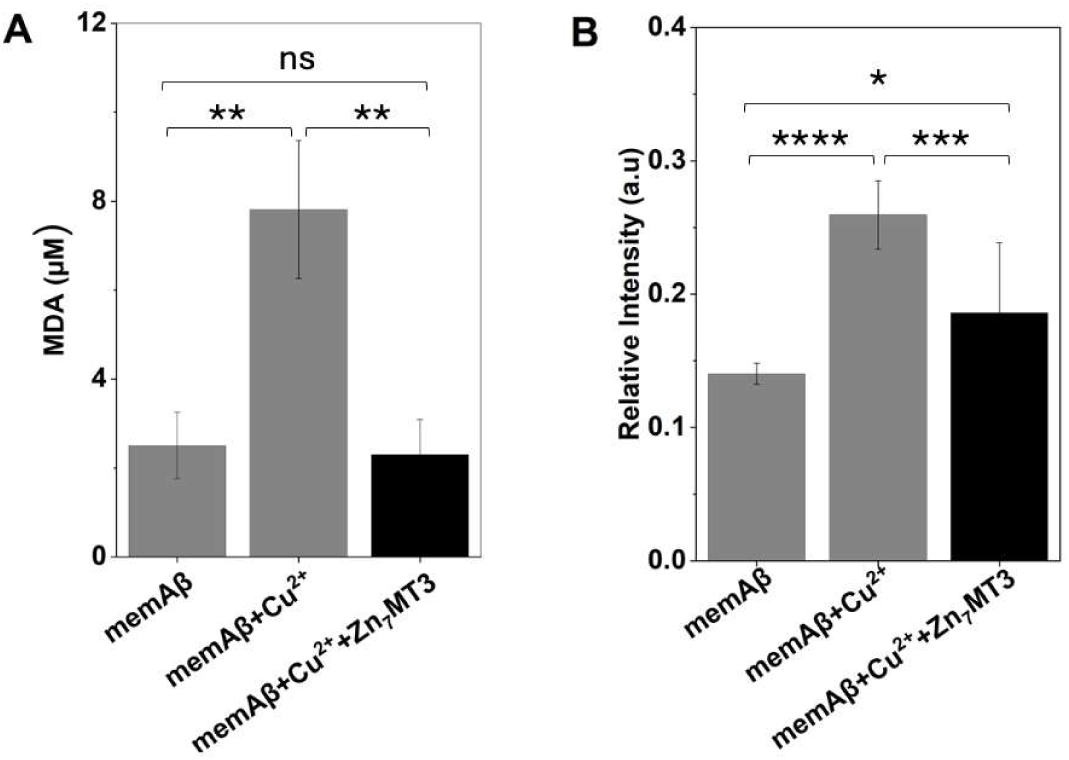
Lipid peroxidation catalyzed by memAβ-Cu^2+^ is silenced by Zn_7_MT-3. A) Quantification of the arachidonic acid (AA) lipid peroxidation product, MDA, in memAβ_1-42_ (12 µM; gray), memAβ_1-42_ -Cu^2+^ (10 µM; black), and upon a metal swap reaction with Zn_7_MT-3 (2.5 µM; black) (n = 4 ± s.d.). B) Membrane leakage quantification by fluorescence upon encapsulation of 5(6)-carboxyfluorescein in the SUV lumen determined for memAβ_1-42_ (12 µM; gray), memAβ_1-42_ -Cu^2+^ (10 µM; black), and upon a metal swap reaction with Zn_7_MT-3 (2.5 µM; black) (n = 9 ± s.d.). Zn_7_MT-3 significantly protects the structural integrity of the membrane. For statistical analysis, a student’s t test was performed (95% confidence): P>0.05 (ns), P≤0.05 (*), P≤10^-2^ (**), P≤10^-3^ (***), P≤10^-4^ (****).

These findings and investigations demonstrate that Zn_7_MT-3 effectively remove Cu^2+^ from membrane-bound Aβ_1-42_-Cu^2+^ as efficiently as it does with soluble Aβ_1-42_-Cu^2+^ complexes, preventing Cu^2+^-mediated ROS production and oxidative cross-linking. By sequestering Cu^+^ in a redox-inactive cluster, MT-3 eliminates memAβ_1-42_-Cu^2+^ pro-oxidant activity, highlighting its central and multi-faceted protective role in mitigating copper-driven membrane-associated neurotoxic processes.

Given its high neuronal expression and distinct copper-selectivity bias for Cu^+^ over other metallothionein isoforms, MT-3 plays a pivotal role in maintaining copper homeostasis within the CNS and regulating aberrant metal-protein interactions.

The role of MT-3 in neurodegenerative diseases (NDs) is underscored by its downregulation in Alzheimer’s disease (AD).^34^ Originally identified as growth inhibitory factor (GIF), which suppresses abnormal neuronal sprouting,^33^ MT-3 was later classified within the metallothionein family.^38^ Unlike the ubiquitously expressed MT-1 and MT-2 isoforms, MT-3 exhibits a unique copper-thionein character, as demonstrated in in-vivo and in-vitro metalation studies. Structural features such as the CPCP motif in the β-domain and the acidic hexapeptide insert in the α-domain enhance its copper selectivity, favoring the formation of more stable Cu^+^ clusters compared to MT-2. With an exceptionally high Cu^+^ affinity (K_d_=10⁻^20^ M), MT-3 functions as a key regulator of copper buffering in the brain, preventing the toxic effects of excess copper.

Dysregulated MT-3 expression has been implicated in several neurodegenerative diseases, including AD, PD, and prion disorders. Given its neuroprotective role in mitigating metal-associated toxicity, strategies to induce MT-3 expression are being explored as potential therapeutic approaches. For example, benzothiazolone-2,^71^ has been shown to upregulate MT-3 with minimal cytotoxicity, while dexamethasone has been observed to suppress Cu^2+^-induced α-Syn aggregation *in vivo* through MT induction.

Our study provides critical mechanistic insight into the role of MT-3 in neutralizing all physiologically relevant Aβ-Cu^2+^ complexes, including the highly toxic membrane-bound forms. By sequestering Cu^2+^ from memAβ-Cu^2+^ and promoting its reduction to Cu^+^, MT-3 prevents the deleterious redox activity, effectively abolishing dopamine oxidation, ROS production, and preventing lipid peroxidation, promoting membrane integrity. This mechanistic framework highlights MT-3 as a key antioxidant and modulator of copper-mediated neurotoxicity and reinforces its potential as a therapeutic target for metal-associated neurodegenerative diseases, in particular AD.

## 4. Conclusion

This works provides novel molecular insights into the role of membrane-bound Aβ-Cu^2+^ complexes in Alzheimer’s disease (AD) pathology, significantly substantiating their contribution to neurotoxicity and oxidative stress in the AD brain. By demonstrating that Aβ_1-42_ can efficiently insert into lipid bilayers without losing its ability to coordinate Cu^2+^, we establish a novel mechanism through which Aβ-Cu interactions can exacerbate neuronal toxicity and damage. The enhanced catalytic catechol oxidase activity towards neurotransmitters like dopamine, ROS production, and the enhancement of lipid peroxidation unraveled for membrane-bound Aβ_1-42_-Cu^2+^ suggest that this species plays a significant role in compromising membrane integrity, ultimately potentially leading to cellular dysfunction and death. This mechanistic understanding underscores the importance of investigating membrane-associated Aβ species as key contributors to AD progression.

This study provides also novel insides on the protective role of metallothionein-3 (MT-3) in preventing the toxic effects of Aβ_1-42_-Cu^2+^. MT-3 can efficiently sequester Cu^2+^ from membrane-bound Aβ species preventing their participation in redox reactions that can generate toxic ROS and drive lipid peroxidation. The ability of MT-3 to undergo a metal swap reaction, reducing Cu^2+^ to Cu^+^ in a unique redox-inert metal-thiolate cluster, constitute a powerful neuroprotective mechanism that could be further explored for therapeutic applications. Given that MT-3 levels are downregulated in AD patients, this finding suggests that boosting MT-3 expression or mimicking its metal-binding properties could be a promising strategy for mitigating also the membrane-associated Aβ-Cu^2+^-induced toxicity.

Overall, this study provides evidence that membrane-bound Aβ-Cu^2+^ complexes are players in AD pathogenesis and that targeting these species could offer new therapeutic opportunities. Future research should focus on developing strategies to enhance MT-3 activity or design biomimetic molecules capable of neutralizing membrane Aβ-Cu^2+^ mediated toxicity. Additionally, further *in vivo* investigations are necessary to confirm the translational potential of these findings and, as well as to explore their implications for AD treatment. By deepening our understanding of the interplay between Aβ, Cu^2+^, and lipid membranes, this work paves the way for novel molecular approaches to combat AD.

## Supporting information

Supplementary Infromation

## Acknowledgements

The work was supported by the National Institute of General Medical Sciences (NIH, R35GM128704 to G.M.), and the Robert A. Welch Foundation (AT-1935-20170325 and AT-2073-20210327 to G.M.). L.P.M. gratefully acknowledges financial support from the National Council of Science and Technology of Mexico (CONACYT) via a doctoral fellowship. We thank Fernando Montalvillo Ortega for generating the membrane-bound Aβ model used in the graphical abstract. We also thank Dr. Mitchell A. Pope, Dr. Jenifer S. Calvo, and Dr. Fabian C. Herbert for helpful discussions. The scheme in **Figure 9** and graphical abstract were created with BioRender.com. CHARMM-GUI and VMD software were utilized for the graphical abstract as well.^72,73^

## Abbreviations

AD: Alzheimer’s Disease
Aβ: Amyloid-beta
APP: Amyloid Precursor Protein
AA: Arachidonic Acid
AALip: Lipids with 1% Arachidonic Acid
CNS: Central Nervous System
GIF: Growth Inhibitory Factor
ICP-MS: Inductively Coupled Plasma-Mass Spectrometry
PD: Parkison’s Disease
MBTH: 3-Methyl-2-Benzothiazoline Hydrazone
MPAC: Metal Protein Attenuating Compounds
MDA: Malondialdehyde
MT: Metallothionein
MT-3: Metallothionein-3
POPC: Phosphatidylcholine
POPG: Phosphatidylglycerol
PrP: Prion Protein
PT: Pellet
ROS: Reactive Oxygen Species
SEC: Size-Exclusion Chromatography
SN: Supernatant Fraction
SUVs: Small Unilamellar Vesicles
THB: 1,2,3-Trihydroxylbenzene
α-Syn: Alpha-synuclein
3-CCA: Coumarin-3-Carboxylic Acid
7-OH-3-CCA: 7-Hydroxycoumarin-3-Carboxylic Acid

