## Supplementary Infromation for "Redox reactivities of membrane-bound Amyloid-β-Cu complexes and their targeting by metallothionein-3"

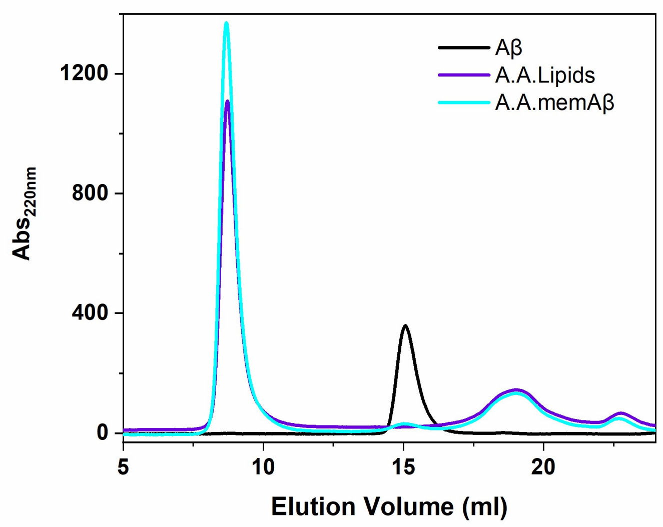


**Supplementary Figure 1**. Size exclusion chromatogram of soluble A$\beta_{1-42}$ (20 µM; black), SUVs (10mM total lipids) doped with 1% arachidonic acid (w/w.; labeled A.A.) (purple), and A$\beta_{1-42}$ incubated with SUVs (10mM total lipid) doped with 1% arachidonic acid (cyan), eluted in 20 mM phosphate buffer pH 7.4.

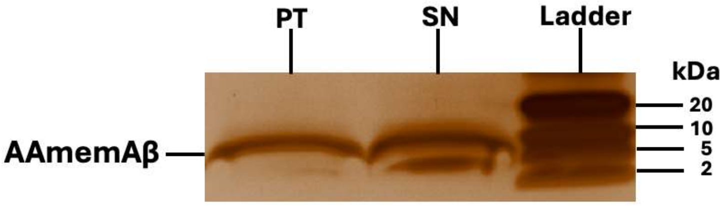


**Supplementary Figure 2**. SDS-PAGE silver staining analysis of a memA$\beta_{1\text{-}42}$-$Cu^{2+}$ in the SUV (with 1% w/w A.A.) pellet (PT) and supernatant (SN) fractions after ultracentrifugation centrifugation in 20 mM phosphate buffer pH 7.4.

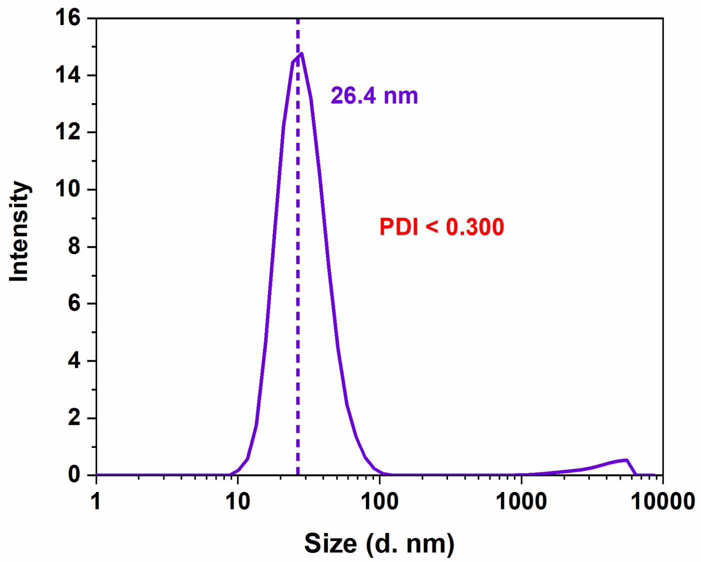


**Supplementary Figure 3**. Dynamic light scattering analysis of SUVs doped with 1% (w/w) arachidonic acid after encapsulation of 5(6)-carboxyfluorescein in the vesicle lumen

| $H_2O_2$ (mM) | $V_{max}$ (nmol Dopaquinone µmol$^{-1}$ Aβ-$Cu^{2+}$ * min$^{-1}$) | $V_{max}$ (nmol Dopaquinone µmol$^{-1}$ memAβ-$Cu^{2+}$ * min$^{-1}$) |
|---|---|---|
| 0 | 10.31 ± 1.20 | 55.62 ± 6.77 |
| 3 | 14.08 ± 0.66 | 92.29 ± 10.34 |
| 8 | 18.58 ± 0.91 | 108.03 ± 8.00 |
| 15 | 16.99 ± 1.04 | 119.61 ± 8.94 |
| 30 | 20.28 ± 1.10 | 112.81 ± 5.22 |
| 50 | 21.11 ± 1.32 | 99.19 ± 6.74 |

**Supplementary Table S1**. Catechol oxidase activity $V_{max}$ values for memAβ-$Cu^{2+}$ and soluble Aβ-$Cu^{2+}$ as a function of $H_2O_2$ concentration.

| | τ (μs) | |
|---|---|---|
| **Sample** | **425 nm** | **575 nm** |
| **memAβ-$Cu^{2+}$ + $Zn_7$MT-3** | 41.2 ± 0.9 | 127.1 ± 2.5 |
| **Aβ-$Cu^{2+}$ + $Zn_7$MT-3** | 42.7 ± 0.5 | 133.1 ± 1.5 |
| **$Cu^{2+}$ + $Zn_7$MT-3** | 42.3 ± 1.0 | 134.7 ± 2.4 |

**Supplementary Table S2**. Low-temperature (77K) luminescence emission lifetimes at 425 and 575 nm for MT-3 upon reaction with memAβ-$Cu^{2+}$, using soluble Aβ-$Cu^{2+}$ species or $Cu^{2+}$ reacted with $Zn_7$MT-3 as references.
